# The neural architecture of language: Integrative modeling converges on predictive processing

**DOI:** 10.1101/2020.06.26.174482

**Authors:** Martin Schrimpf, Idan Blank, Greta Tuckute, Carina Kauf, Eghbal A. Hosseini, Nancy Kanwisher, Joshua Tenenbaum, Evelina Fedorenko

## Abstract

The neuroscience of perception has recently been revolutionized with an integrative modeling approach in which computation, brain function, and behavior are linked across many datasets and many computational models. By revealing trends across models, this approach yields novel insights into cognitive and neural mechanisms in the target domain. We here present a first systematic study taking this approach to higher-level cognition: human language processing, our species’ signature cognitive skill. We find that the most powerful ‘transformer’ models predict nearly 100% of explainable variance in neural responses to sentences and generalize across different datasets and imaging modalities (fMRI, ECoG). Models’ neural fits (‘brain score’) and fits to behavioral responses are both strongly correlated with model accuracy on the next-word prediction task (but not other language tasks). Model architecture appears to substantially contribute to neural fit. These results provide computationally explicit evidence that predictive processing fundamentally shapes the language comprehension mechanisms in the human brain.

**Significance:** Language is a quintessentially human ability. Research has long probed the functional architecture of language processing in the mind and brain using diverse brain imaging, behavioral, and computational modeling approaches. However, adequate neurally mechanistic accounts of how meaning might be extracted from language are sorely lacking. Here, we report an important first step toward addressing this gap by connecting recent artificial neural networks from machine learning to human recordings during language processing. We find that the most powerful models predict neural and behavioral responses across different datasets up to noise levels. Models that perform better at predicting the next word in a sequence also better predict brain measurements – providing computationally explicit evidence that predictive processing fundamentally shapes the language comprehension mechanisms in the human brain.

---

A core goal of neuroscience is to decipher from patterns of neural activity the algorithms underlying our abilities to perceive, think, and act. Recently, a new “reverse engineering” approach to computational modeling in systems neuroscience has transformed our algorithmic understanding of the primate ventral visual stream (Bao et al., 2020; Cadena et al., 2019; Cichy et al., 2016; Kietzmann et al., 2019; Kubilius et al., 2019; Schrimpf et al., 2018, 2020; Yamins et al., 2014), and holds great promise for other aspects of brain function. This approach has been enabled by a breakthrough in artificial intelligence (AI): the engineering of artificial neural network (ANN) systems that perform core perceptual tasks with unprecedented accuracy, approaching human levels, and that do so using computational machinery that is abstractly similar to biological neurons. In the ventral visual stream, the key AI developments come from deep convolutional neural networks (DCNNs) that perform visual object recognition from natural images (Cireşan et al., 2012; Krizhevsky et al., 2012; Schrimpf et al., 2018, 2020; Yamins et al., 2014), widely thought to be the primary function of this pathway. Leading DCNNs for object recognition have now been shown to predict the responses of neural populations in multiple stages of the ventral stream (V1, V2, V4, IT), in both macaque and human brains, approaching the noise ceiling of the data. Thus, despite abstracting away aspects of biology, DCNNs provide the basis for a first complete hypothesis of how the brain extracts object percepts from visual input.

Inspired by this success story, analogous ANN models have now been applied to other domains of perception (Kell et al., 2018; Zhuang et al., 2017). Could these models also let us reverse-engineer the brain mechanisms of higher-level human cognition? Here we show for the first time how the modeling approach pioneered in the ventral stream can be applied to a higher-level cognitive domain that plays an essential role in human life: language comprehension, or the extraction of meaning from spoken, written or signed words and sentences. Cognitive scientists have long treated neural network models of language processing with skepticism (Marcus, 2018; Pinker & Prince, 1988) given that these systems lack (and often deliberately attempt to do without) explicit symbolic representation – traditionally seen as a core feature of linguistic meaning. Recent ANN models of language, however, have proven capable of at least approximating some aspects of symbolic computation, and have achieved remarkable success on a wide range of applied natural language processing (NLP) tasks. The results presented here, based on this new generation of ANNs, suggest that a computationally adequate model of language processing in the brain may be closer than previously thought.

Because we build on the same logic in our analysis of language in the brain, it is helpful to review why the neural network-based integrative modeling approach has proven so powerful in the study of object recognition in the ventral stream. Crucially, our ability to robustly link computation, brain function, and behavior is supported not by testing a single model on a single dataset or a single kind of data, but by large-scale *integrative benchmarking* (Schrimpf et al., 2020) that establishes consistent patterns of performance across many different ANNs applied to multiple neural and behavioral datasets, together with their performance on the proposed core computational function of the brain system under study. Given the complexities of the brain’s structure and the functions it performs, any one of these models is surely oversimplified and ultimately wrong – at best, an approximation of some aspects of what the brain does. But some models are less wrong than others, and consistent *trends in performance across models* can reveal not just which model best fits the brain, but which properties of a model underlie its fit to the brain, thus yielding critical insights that transcend what any single model can tell us.

In the ventral stream specifically, our understanding that computations underlying object recognition are analogous to the structure and function of DCNNs is supported by findings that across hundreds of model variants, DCNNs that perform better on object recognition tasks also better capture human recognition behavior and neural responses in IT cortex of both human and non-human primates (Rajalingham et al., 2018; Schrimpf et al., 2018, 2020; Yamins et al., 2014). This integrative benchmarking reveals a rich pattern of correlations among three classes of performance measures — (i) neural variance explained, in IT neurophysiology or fMRI responses (brain scores), (ii) accuracy in predicting hits and misses in human object recognition behavior, or human object similarity judgments (behavioral scores), and (iii) accuracy on the core object recognition task (computational task score) — such that for any individual DCNN model we can predict how well it would score on each of these measures from the other measures. This pattern of results was not assembled in a single paper but in multiple papers across several labs and several years. Taken together, they provide strong evidence that the ventral stream supports primate object recognition through something like a deep convolutional feature hierarchy, the exact details of which are being modeled with ever-increasing precision.

Here we describe an analogous pattern of results for ANN models of human language, establishing a link between language models, including transformer-based ANN architectures that have revolutionized natural language processing in AI systems over the last three years, and fundamental computations of human language processing as reflected in both neural and behavioral measures. Language processing is known to depend causally on a left-lateralized fronto-temporal brain network (Bates et al., 2003; Binder et al., 1997; Fedorenko & Thompson-Schill, 2014; Friederici, 2012; Gorno-Tempini et al., 2004; Hagoort, 2019; Price, 2010) (Fig. 1) that responds robustly and selectively to linguistic input (Fedorenko et al., 2011; Monti et al., 2012), whether auditory or visual (Deniz et al., 2019; Regev et al., 2013). Yet the precise computations underlying language processing in the brain remain unknown. Computational models of sentence processing have previously been used to explain both behavioral (Dotlačil, 2018; Futrell, Gibson, & Levy, 2020; Gibson, 1998; Gibson et al., 2013; Hale, 2001; Jurafsky, 1996; Lakretz et al., 2020; Levy, 2008a, 2008b; Lewis et al., 2006; McDonald & Macwhinney, 1998; Smith & Levy, 2013; Spivey-Knowlton, 1996; Steedman, 2000; van Schijndel et al., 2013), and neural responses to linguistic input (Brennan et al., 2016; Brennan & Pylkkänen, 2017; Ding et al., 2015; Frank et al., 2015; Henderson et al., 2016; Huth et al., 2016; Lopopolo et al., 2017; Lyu et al., 2019; T. M. Mitchell et al., 2008; Nelson et al., 2017; Pallier et al., 2011; Pereira et al., 2018; Rabovsky et al., 2018; Shain et al., 2020; Wehbe et al., 2014; Willems et al., 2016; Gauthier & Ivanova, 2018; Gauthier & Levy, 2019; Hu et al., 2020; Jain & Huth, 2018; S. Wang et al., 2020; Schwartz et al., 2019; Toneva & Wehbe, 2019). However, none of the prior studies have attempted large-scale integrative benchmarking that has proven so valuable in understanding key brain-behavior-computation relationships in the ventral stream; instead, they have typically tested one or a small number of models against a single dataset, and the same models have not been evaluated on all three metrics of neural, behavioral, and objective task performance. Previously tested models have also left much of the variance in human neural/behavioral data unexplained. Finally, until the rise of recent ANNs (e.g., transformer architectures), language models did not have sufficient capacity to solve the full linguistic problem that the brain solves – to form a representation of sentence meaning capable of performing a broad range of real-world language tasks on diverse natural linguistic input. We are thus left with a collection of suggestive results but no clear sense of how close ANN models are to fully explaining language processing in the brain, or what model features are key in enabling models to explain neural and behavioral data.

**Figure 1:**
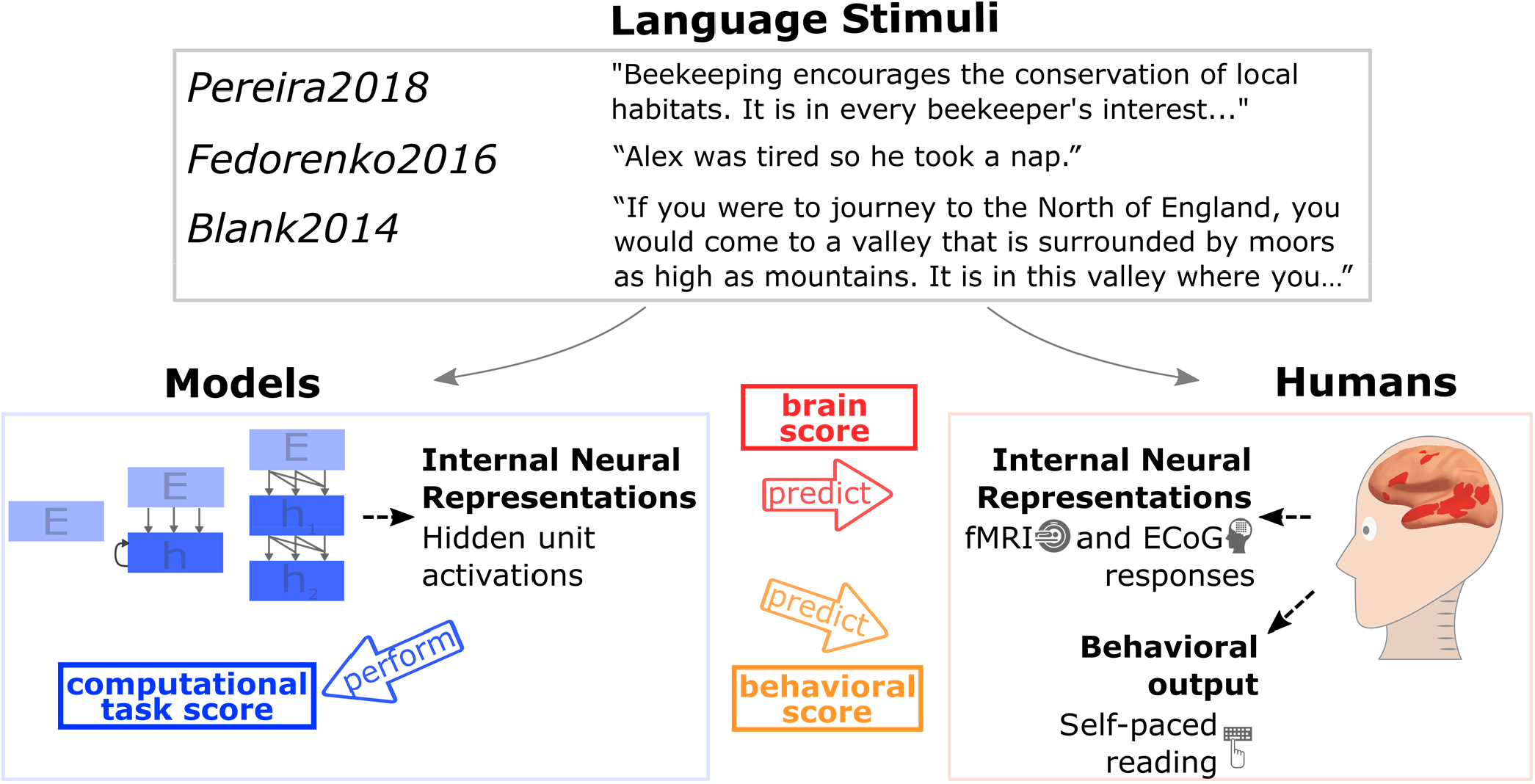
Comparing Artificial Neural Network models of language processing to human language processing. We tested how well different models predict measurements of human neural activity (fMRI and ECoG) and behavior (reading times) during language comprehension. The candidate models ranged from simple embedding models to more complex recurrent and transformer networks. Stimuli ranged from sentences to passages to stories and were 1) fed into the models, and 2) presented to human participants (visually or auditorily). Models’ internal representations were evaluated on three major dimensions: their ability to predict human neural representations (brain score, extracted from within the fronto-temporal language network (e.g., Fedorenko et al., 2010; the network topography is schematically illustrated in red on the template brain above); their ability to predict human behavior in the form of reading times (behavioral score); and their ability to perform computational tasks such as next-word prediction (computational task score). Consistent relationships between these measures across many different models reveal insights beyond what a single model can tell us.

Our goal here is to present a first systematic integrative modeling study of language in the brain, at the scale necessary to discover robust relationships between neural and behavioral measurements from humans, and performance of models on language tasks. We seek to determine not just which model fits empirical data best, but what dimensions of variation across models are correlated with fit to human data. This approach has not been applied in the study of language or any other higher cognitive system, and even in perception has not been attempted within a single integrated study. Thus, we view our work more generally as a *template for how to apply the integrative benchmarking approach to any perceptual or cognitive system*.

Specifically, we examined the relationships between 43 diverse state-of-the-art ANN language models (henceforth ’models’) across three neural language comprehension datasets (two fMRI, one electrocorticography (ECoG)), as well as behavioral signatures of human language processing in the form of self-paced reading times, and a range of linguistic functions assessed via standard engineering tasks from NLP. The models spanned all major classes of existing ANN language approaches and included simple embedding models (e.g., GloVe (Pennington et al., 2014)), more complex recurrent neural networks (e.g., LM1B (Jozefowicz et al., 2016)), and many variants of transformers or attention-based architectures—including both ‘unidirectional-attention’ models (trained to predict the next word given the previous words; e.g., GPT (Radford et al., 2019)) and ‘bidirectional-attention’ models (trained to predict a missing word given the surrounding context; e.g., BERT (Devlin et al., 2018)).

Our integrative approach yielded four major findings. (1) Models’ relative fit to neural data (neural predictivity or “brain score’’)—estimated on held-out test data—generalizes across different datasets and imaging modality (fMRI, ECoG), and certain architectural features consistently lead to more brain-like models: transformer-based models perform better than recurrent networks or word-level embedding models, and larger-capacity models perform better than smaller models. (2) The best models explain nearly 100% of the explainable variance (up to the noise ceiling) in neural responses to sentences. This result stands in stark contrast to earlier generations of models that have typically accounted for at most 30-50% of the predictable neural signal. (3) Across models, significant correlations hold among all three metrics of model performance: brain scores (fit to fMRI and ECoG data), behavioral scores (fit to reading time), and model accuracy on the next-word prediction task. Importantly, no other linguistic task was predictive of models’ fit to neural or behavioral data. These findings provide strong evidence for a classic hypothesis about the computations underlying human language understanding, that the brain’s language system is optimized for predictive processing in the service of meaning extraction. (4) Intriguingly, the scores of models initialized with random weights (prior to training, but with a trained linear readout) are well above chance and correlate with trained model scores, which suggests that network architecture is an important contributor to a model’s brain score. In particular, one architecture introduced just in 2019, the generative pre-trained transformer (GPT-2), consistently outperforms all other models and explains almost all variance in both fMRI and ECoG data from sentence processing tasks. GPT-2 is also arguably the most cognitively plausible of the transformer models (because it uses unidirectional, forward attention), and performs best overall as an AI system when considering both natural language understanding and natural language generation tasks. Thus, even though the goal of contemporary AI is to improve model performance and not necessarily to build models of brain processing, this endeavor appears to be rapidly converging on architectures that might capture key aspects of language processing in the human mind and brain.

## Results

We evaluated a broad range of state-of-the-art ANN language models on the match of their internal representations to three human neural datasets. The models spanned all major classes of existing language models (<u>Methods_5</u>, Table S11). The neural datasets consisted of i) fMRI activations while participants read short passages, presented one sentence at a time (across two experiments) that spanned diverse topics (*Pereira2018* dataset (Pereira et al., 2018)); ii) ECoG recordings while participants read semantically and syntactically diverse sentences, presented one word at a time (*Fedorenko2016* dataset (Fedorenko et al., 2016)); and iii) fMRI BOLD signal time-series elicited while participants listened to ∼5-minutes-long naturalistic stories (*Blank2014* dataset (Blank et al., 2014)) (<u>Methods_1-3</u>). Thus, the datasets varied in the imaging modality (fMRI/ECoG), the nature of the materials (unconnected sentences/passages/stories), the grain of linguistic units to which responses were recorded (sentences/words/2s-long story fragments), and presentation modality (reading/listening). In most analyses, we consider the overall results across the three neural datasets; when considering the results for the individual neural datasets, we give the most weight to *Pereira2018* because it includes multiple repetitions per stimulus (sentence) within each participant and quantitatively exhibits the highest internal reliability (Fig. S1). Because our research questions concern language processing, we extracted neural responses from language-selective voxels or electrodes that were functionally identified by an extensively validated independent ’localizer‘ task that contrasts reading sentences versus nonword sequences (Fedorenko et al., 2010). This localizer robustly identifies the fronto-temporal language-selective network (<u>Methods_1-3</u>).

To compare a given model to a given dataset, we presented the same stimuli to the model that were presented to humans in neural recording experiments and ‘recorded’ the model’s internal activations (<u>Methods_5-6,</u> Fig. 1). We then tested how well the model recordings could predict the neural recordings for the same stimuli, using a method originally developed for studying visual object recognition (Schrimpf et al., 2018; Yamins et al., 2014). Specifically, using a subset of the stimuli, we fit a linear regression from the model activations to the corresponding human measurements, modeling the response of each voxel (*Pereira2018*) / electrode (*Fedorenko2016*) / brain region (*Blank2014*) as a linear weighted sum of responses of different units from the model. We then computed model predictions by applying the learned regression weights to model activations for the held-out stimuli, and evaluated how well those predictions matched the corresponding held-out human measurements by computing Pearson’s correlation coefficient. We further normalized these correlations by the extrapolated reliability of the particular dataset, which places an upper bound (‘ceiling‘) on the correlation between the neural measurements and any external predictor (<u>Methods_7</u>, Fig. S1). The final measure of a model’s performance (‘score’) on a dataset is thus Pearson’s correlation between model predictions and neural recordings divided by the estimated ceiling and averaged across voxels/electrodes/regions and participants. We report the score for the best-performing layer of each model (<u>Methods_6</u>, Fig. S12) but controlled for the generality of the layer choice in a train/test split (Fig. S2b, c).

### Specific models accurately predict human brain activity

We found (Fig. 2a-b) that specific models predict *Pereira2018* and *Fedorenko2016* datasets with up to 100% predictivity relative to the noise ceiling (<u>Methods_7</u>, Fig. S1). These scores generalize to another metric, “RDM”, based on representational similarity without any fitting (Fig. S2a). The *Blank2014* dataset is also reliably predicted, but with lower predictivity. Models vary substantially in their ability to predict neural data. Generally, embedding models such as GloVe do not perform well on any dataset. In contrast, recurrent networks such as skip-thoughts, as well as transformers such as BERT, predict large portions of the data. The model that predicts the human data best across datasets is GPT2-xl, a unidirectional-attention transformer model, which predicts *Pereira2018* and *Fedorenko2016* at close to 100% of the noise ceiling and is among the highest-performing models on *Blank2014* with 32% normalized predictivity. These scores are higher in the language network than other parts of the brain (SI-4). Intermediate layer representations in the models are most predictive, significantly outperforming representations at the first and output layers (Figs. 2c, S13).

**Figure 2:**
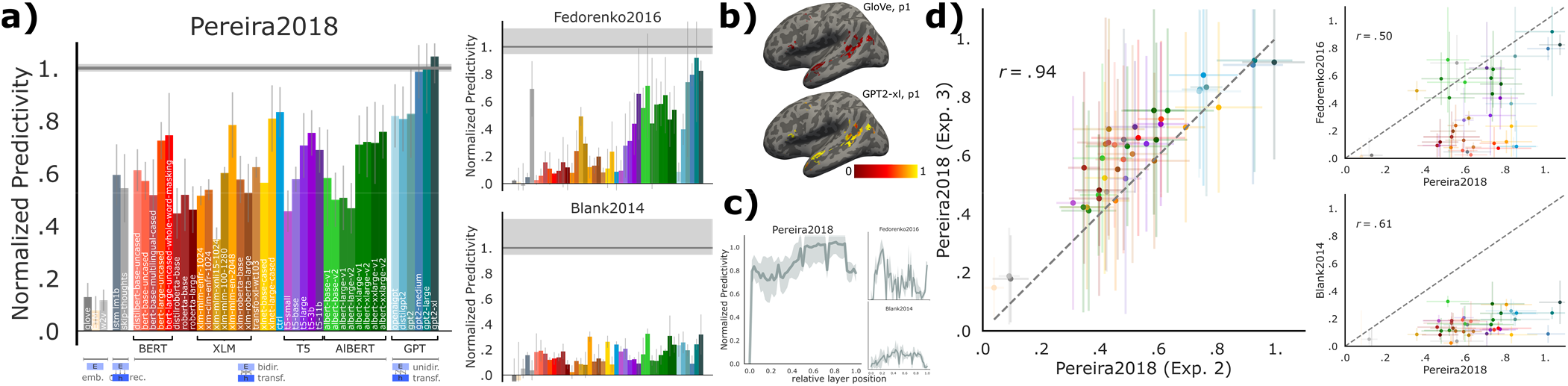
Specific models accurately predict neural responses consistently across datasets. **(a)** We compared 43 computational models of language processing (ranging from embedding to recurrent and bi- and uni-directional transformer models) in their ability to predict human brain data. The neural datasets include: fMRI voxel responses to visually presented (sentence-by-sentence) passages (*Pereira2018*), ECoG electrode responses to visually presented (word-by-word) sentences (*Fedorenko2016*), fMRI region of interest (ROI) responses to auditorily presented ∼5min-long stories (*Blank2014*). For each model, we plot the normalized predictivity (‘brain score’), i.e. the fraction of ceiling (gray line; <u>Methods_7</u>, Fig. S1) that the model can predict. Ceiling levels are .32 (*Pereira2018*), .17 (*Fedorenko2016*), and .20 (*Blank2014*). Model classes are grouped by color (<u>Methods_5</u>, Table S10). Error bars (here and elsewhere) represent median absolute deviation over subject scores. **(b)** Normalized predictivity of GloVe (a low-performing embedding model) and GPT2-xl (a high-performing transformer model) in the language-responsive voxels in the left hemisphere of two representative participants from *Pereira2018* (also Fig. S3). **(c)** Brain score per layer in GPT2-xl. Middle-to-late layers generally yield the highest scores for Pereira2018 and Blank2014 whereas earlier layers better predict Fedorenko2016. This difference might be due to predicting individual word representations (within a sentence) in *Fedorenko2016*, as opposed to whole-sentence representations in *Pereira2018*. **(d)** To test how well model brain scores generalize across datasets, we correlated i) two experiments with different stimuli (and some participant overlap) in *Pereira2018* (obtaining a very strong correlation), an ii) *Pereira2018* brain scores with the scores for each of *Fedorenko2016* and *Blank2014* (obtaining lower but still highly significant correlations). Brain scores thus tend to generalize across datasets, although differences between datasets exist which warrant the full suite of datasets.

### Model scores are consistent across experiments/datasets

To test the generality of the model representations, we examined the consistency of model brain scores across datasets. Indeed, if a model achieves a high brain score on one dataset, it tends to also do well on other datasets (Fig. 2d), ruling out the possibility that we are picking up on spurious, dataset-idiosyncratic predictivity, and suggesting that the models’ internal representations are general enough to capture brain responses to diverse linguistic materials presented visually or auditorily, and across three independent sets of participants. Specifically, model brain scores across the two experiments in *Pereira2018* (overlapping sets of participants) correlate at *r*=.94 (Pearson here and elsewhere, *p<<*.*00001*), scores from *Pereira2018* and *Fedorenko2016* correlate at *r*=.50 (*p<*.*001*), and from *Pereira2018* and *Blank2014* at *r*=.63 (*p<*.*0001*).

### Next-word-prediction task performance selectively predicts brain scores

In the critical test of which computations might underlie human language understanding, we examined the relationship between the models’ ability to predict an upcoming word and their brain scores. Words from the Wikitext-2 dataset (Merity et al., 2016) were sequentially fed into the candidate models. We then fit a linear classifier (over words in the vocabulary; n=50k) from the last layer’s feature representation (frozen, i.e. no finetuning) on the training set to predict the next word, and evaluated performance on the held-out test set (<u>Methods_8</u>). Indeed, next-word-prediction task performance robustly predicts brain scores (Fig. 3a; *r*=.44, *p<*.01, averaged across datasets). The best language model, GPT2-xl, also achieves the highest brain score (see previous section). This relationship holds for model variants within each model class—embedding models, recurrent networks, and transformers— ruling out the possibility that this correlation is due to between-class differences in next-word-prediction performance.

**Figure 3:**
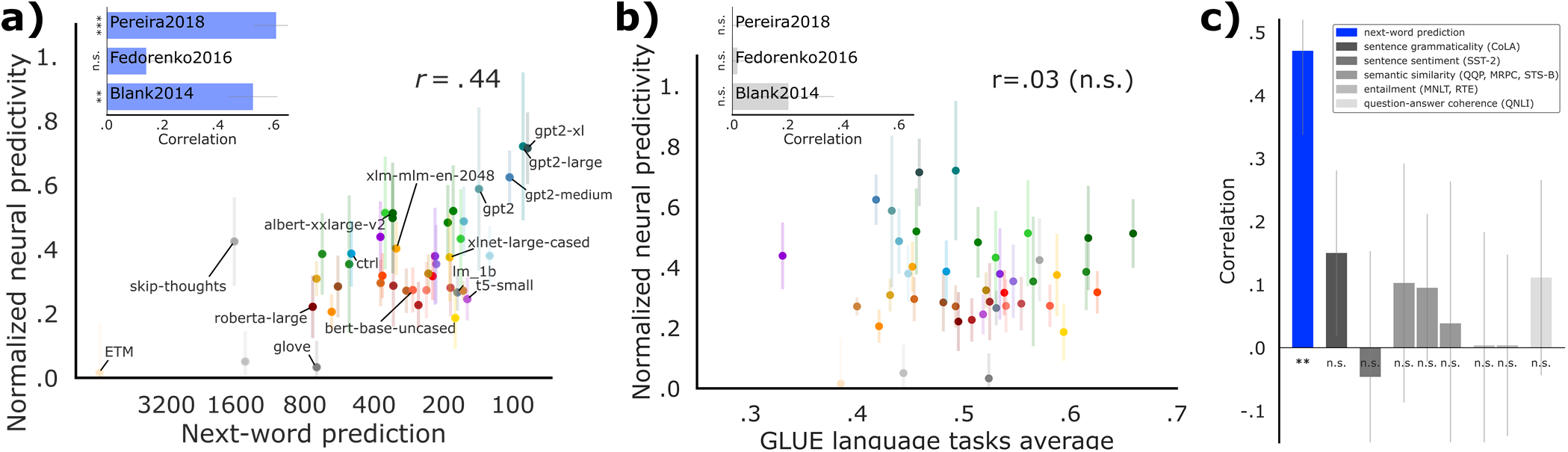
Model performance on a next-word-prediction task selectively predicts brain scores. **(a)** Next-word-prediction task performance was evaluated as the surprisal between the predicted and true next word in the WikiText-2 dataset of 720 Wikipedia articles, or *perplexity* (x-axis, lower is better; training only a linear readout leading to worse perplexity values than canonical fine-tuning, see <u>Methods-8</u>). Next-word-prediction task scores strongly predict brain scores across datasets (inset: this correlation is significant for two individual datasets: *Pereira2018* and *Blank2014*; the correlation for *Fedorenko2016* is positive but not significant). **(b)** Performance on diverse language tasks from the GLUE benchmark collection does *not* correlate with overall or individual-dataset brain scores (inset; SI-5; training only a linear readout). **(c)** Correlations of individual tasks with brain scores. Only improvements on next-word prediction lead to improved neural predictivity.

To test whether next-word prediction is special in this respect, we asked whether model performance on *any* language task correlates with brain scores. As with next-word prediction, we kept the model weights fixed and only trained a linear readout. We found that performance on tasks from the GLUE benchmark collection (Cer et al., 2018; Dolan & Brockett, 2005; Levesque et al., 2012; Rajpurkar et al., 2016; Socher et al., 2013; A. Wang, Singh, et al., 2019; Warstadt et al., 2019; Williams et al., 2018)—including grammaticality judgments, sentence similarity judgments, and entailment—does *not* predict brain scores (Fig. 3b-c). The difference in the strength of correlation between brain scores and the next-word prediction task performance vs. the GLUE tasks performance is highly reliable (p<<0.00001, t-test over 1,000 bootstraps of scores and corresponding correlations; <u>Methods_9</u>). This result suggests that optimizing for predictive representations may be a critical shared objective of biological and artificial neural networks for language, and perhaps more generally (Keller and Mrsic-Flogel, 2018; Singer et al., 2018).

### Brain scores and next-word-prediction task performance correlate with behavioral scores

Beyond internal neural representations, we tested the models’ ability to predict external behavioral outputs because, ultimately, in integrative benchmarking, we strive for a computationally precise account of language processing that can explain both neural response patterns and observable linguistic behaviors. We chose a large corpus (n=180 participants) of self-paced reading times for naturalistic story materials (*Futrell2018* dataset (Futrell, Gibson, Tily, et al., 2020)). Per-word reading times provide a theory-neutral measure of incremental comprehension difficulty, which has long been a cornerstone of psycholinguistic research in testing theories of sentence comprehension (Demberg & Keller, 2008; Gibson, 1998; Just & Carpenter, 1980; D. C. Mitchell, 1984; Rayner, 1978; Smith & Levy, 2013) and which were recently shown to robustly predict neural activity in the language network (Wehbe et al., 2020).

### Specific models accurately predict reading times

We regressed each model’s last layer’s feature representation (i.e., closest to the output) against reading times and evaluated predictivity on held-out words. As with the neural datasets, we observed a spread of model ability to capture human behavioral data, with models such as GPT2-xl and AlBERT-xxlarge predicting these data close to the noise ceiling (Fig. 4a; also Merkx & Frank, 2020; Wilcox et al., 2020).

**Figure 4:**
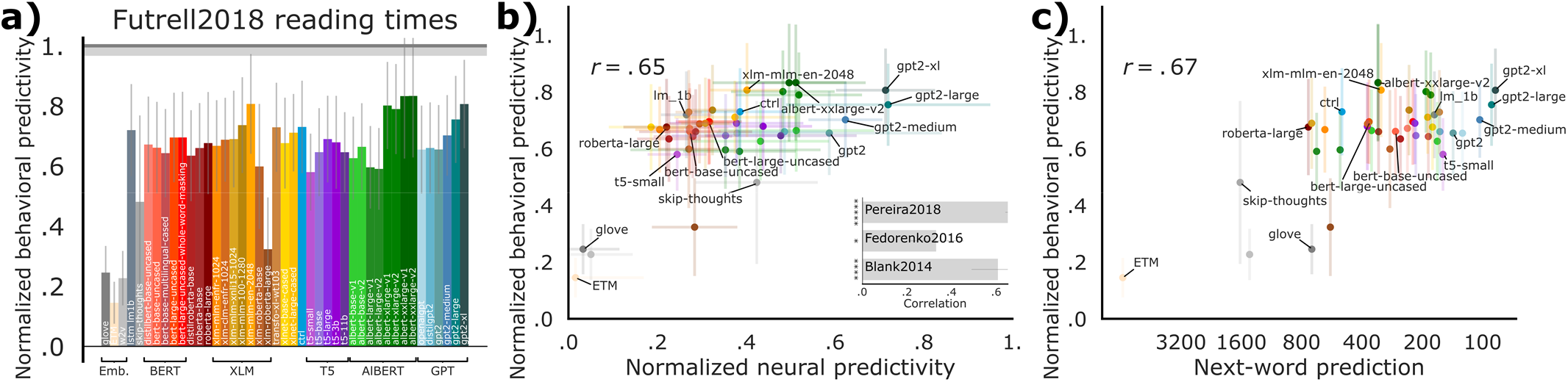
Behavioral scores, brain scores, and next-word-prediction task performance are pairwise correlated. **(a)** Behavioral predictivity of each model on *Futrell2018* reading times (notation similar to Fig. 2). Ceiling level is .76. **(b)** Models’ neural scores aggregated across the three neural datasets (or for each dataset individiually; inset and Fig. S6) correlates with behavioral scores. **(c)** Next-word-prediction task performance (Fig. 3) correlates with behavioral scores. Performance on other language tasks (from the GLUE benchmark collection) does *not* correlate with behavioral scores (Fig. S7).

### Brain scores correlate with behavioral scores

To test whether models with the highest brain scores also predict reading times best, we compared models’ neural predictivity (across datasets) with those same models’ behavioral predictivity. Indeed, we observed a strong correlation (Fig. 4b; r=.65, *p<<*.*0001*), which also holds for the individual neural datasets (inset and Fig. S6). These results suggest that further improving models’ neural predictivity will simultaneously improve their behavioral predictivity.

### Next-word-prediction task performance correlates with behavioral scores

Next-word-prediction task performance is predictive of reading times (Fig. 4c; r=.67, *p<<*.*0001)*, in line with earlier studies (Goodkind & Bicknell, 2018; van Schijndel & Linzen, 2018) and thus connecting all three measures of performance: brain scores, behavioral scores, and task performance on next-word prediction.

### Model architecture contributes to model-to-brain relationship

The brain’s language network plausibly arises through a combination of evolutionary and learning-based optimization. In a first attempt to test the relative importance of the models’ intrinsic architectural properties vs. training-related features, we performed two analyses. First, we found that architectural features (e.g. number of layers) but neither of the features related to training (e.g. dataset and vocabulary size) significantly predicted improvements in model performance on the neural data (S10, Table S11). These results align with prior studies that had reported that architectural differences affect model performance on normative tasks like next-word prediction after training, and define the representational space that the model can learn (Arora et al., 2018; Fukushima, 1988; Geiger et al., 2020). Second, we computed brain scores for the 43 models without training, i.e. with initial (random) weights. Note that the predictivity metric still trains a linear readout on top of the model representations. Surprisingly, even with no training, several models achieved reasonable scores (Fig. 5), consistent with recent results of models in high-level visual cortex (Geiger et al., 2020) as well as findings on the power of random initializations in natural language processing (Merchant et al., 2020; Tenney et al., 2019; Zhang & Bowman, 2018). For example, across the three datasets, untrained GPT2-xl achieves an average predictivity of ∼*51%*, only ∼*20%* lower than the trained network. A similar trend is observed across models: training generally improves brain scores, on average by 53%. Across models, the untrained scores are strongly predictive of the trained scores (*r*=.74, *p<<*.*00001*), indicating that models that already perform well with random weights improve further with training.

**Figure 5:**
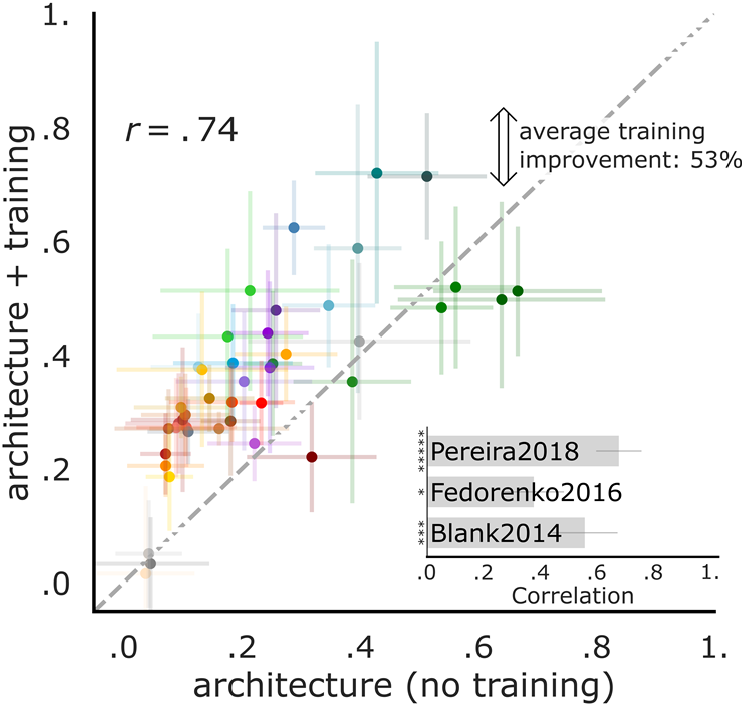
Model architecture contributes to the model-brain relationship. We evaluate untrained models by keeping weights at their initial random values. The remaining representations are driven by architecture alone and are tested on the neural datasets (Fig. 2). Across the three datasets, architecture alone yields representations that predict human brain activity considerably well. On average, training improves model scores by 53%. For *Pereira2018*, training improves predictivity the most whereas for *Fedorenko2016* and *Blank2014*, training does not always change—and for some models even decreases—neural scores (Fig. S8). The untrained model performance is consistently predictive of its performance after training across and within (inset) datasets.

To ensure the robustness and generalizability of the results for untrained models, and to gain further insights into these results, we performed four additional analyses (Fig. S9). First, we tested a random context-independent embedding with equal dimensionality to the GPT2-xl model but no architectural priors and found that it predicts only a small fraction of the neural data, on average below 15%, suggesting that a large feature space alone is not sufficient (Fig. S9a). Second, to ensure that the overlap between the linguistic materials (words, bigrams, etc.) used in the train and test splits is not driving the results, we quantified the overlap and found it to be low, especially for bi- and tri-grams (Fig. S9b). Third, to ensure that the linear regression used in the predictivity metric did not artificially inflate the scores of untrained models, we used an alternative metric – “RDM” – that does not involve any fitting. Scores of untrained models on the predictivity metric generalized to scores on the RDM metric (Fig. S9d). Finally, we examined the performance of untrained models with a trained linear readout on the next-word prediction task and found similar performance trends to those we observed for the neural scores (Fig. S9c), confirming the representational power of untrained representations.

## Discussion

### Summary of key results and their implications

Our results, summarized in Fig. 6, show that specific ANN language models can predict human neural and behavioral responses to linguistic input with high accuracy: the best models achieve, on some datasets, perfect predictivity relative to the noise ceiling. Model scores correlate across neural and behavioral datasets spanning recording modalities (fMRI, ECoG, reading times) and diverse materials presented visually and auditorily across three sets of participants, establishing the robustness and generality of these findings. Critically, both neural and behavioral scores correlate with model performance on the normative next-word prediction task – but not other language tasks. Finally, untrained models with random weights (and a trained linear readout) produce representations beginning to approximate those in the brain’s language network.

**Figure 6 (Overview of results):**
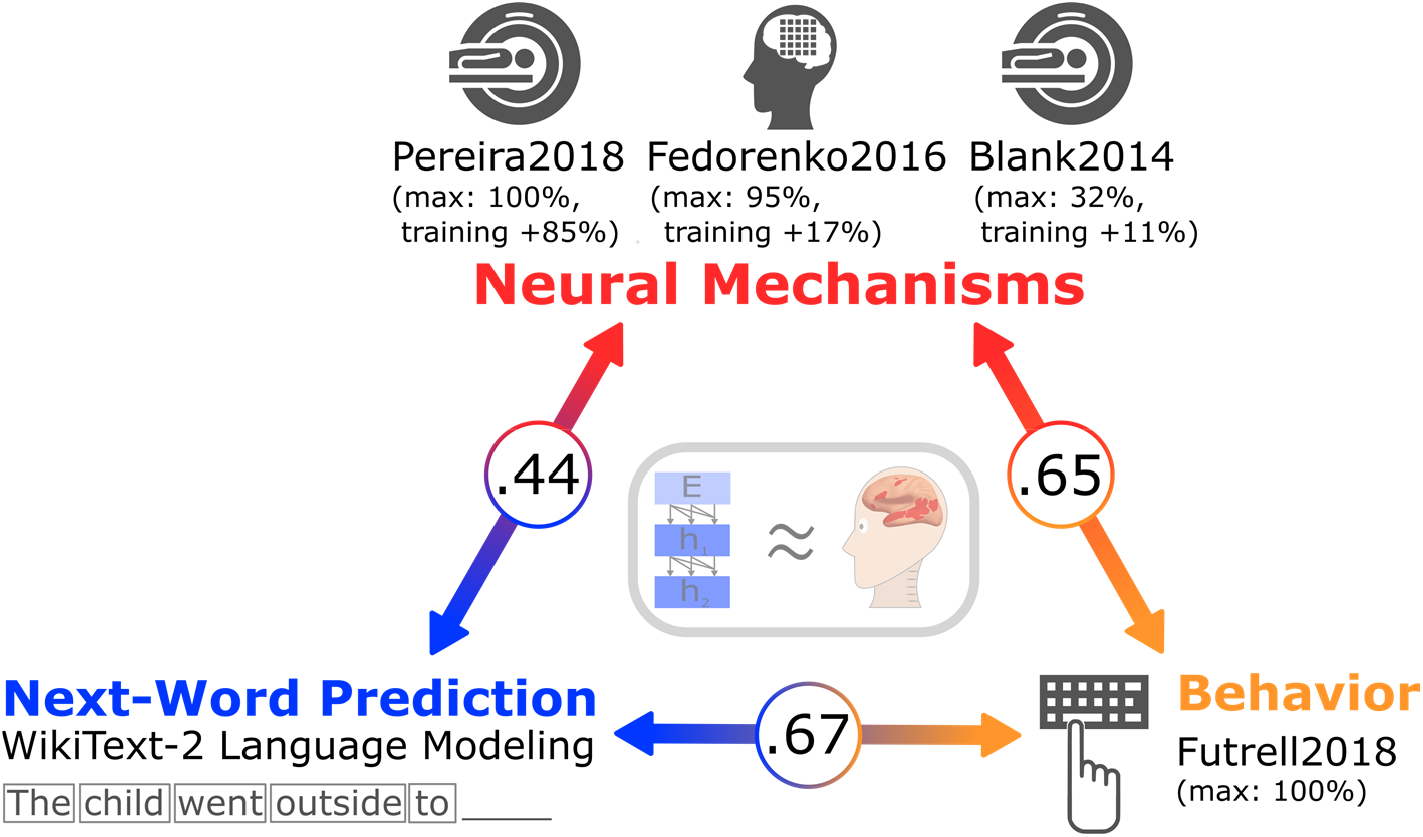
Connecting neural mechanisms, behavior, and computational task (next-word prediction). Specific ANN language models are beginning to approximate the brain’s mechanisms for processing language (middle gray box). For the neural datasets (fMRI and ECoG recordings; top, red), and for the behavioral dataset (self-paced reading times; bottom right, orange), we report i) the value for the model achieving the highest predictivity, and ii) the average improvement on brain scores across models after training. Model performances on the next-word-prediction task (WikiText-2 language modeling perplexity; bottom left, blue) predict brain and behavioral scores; and brain scores predict behavioral scores (circled numbers).

### Predictive language processing

Underlying the integrative modeling framework, implemented here in the cognitive domain of language, is the idea that large-scale neural networks can serve as hypotheses of the actual computations conducted in the brain. We here identified some models—unidirectional-attention transformer architectures—that accurately capture brain activity during language processing. We then began dissecting variations across the range of model candidates to explain *why* they achieve high brain scores. Two core findings emerged, both supporting the idea that the human language system is optimized for predictive processing. First, we found that the models’ performance on the next-word prediction task, but not other language tasks, is correlated with neural predictivity (see (Gauthier & Levy, 2019) for related evidence of fine-tuning of one model on tasks other than next-word-prediction leading to worse model-to-brain fit). Recent preprints conceptually replicate and extend this basic finding (Caucheteux & King, 2020; Goldstein et al., 2020; Wehbe et al., 2020; Wilcox et al., 2020). Language modeling (predicting the next word) is the task of choice in the natural language processing (NLP) community: it is simple, unsupervised, scalable, and appears to produce the most generally useful, successful language representations. This is likely because language modeling encourages a neural network to build a joint probability model of the linguistic signal, which implicitly requires sensitivity to diverse kinds of regularities in the signal.

Second, we found that the models that best match human language processing are precisely those that are trained to predict the next word. Predictive processing has advanced to the forefront of theorizing in cognitive science (Christiansen & Chater, 1999; Clark, 2013; Elman, 1990, 1991, 1993; McRae et al., 1998; Rohde & Plaut, 1999; Spivey & Tanenhaus, 1998; Tenenbaum et al., 2011) and neuroscience (Bastos et al., 2012; Keller & Mrsic-Flogel, 2018; Mumford, 1992; Rao & Ballard, 1999; Srinivasan et al., 1982), including in the domain of language (Kuperberg & Jaeger, 2016; Levy, 2008a). The rich sources of information that comprehenders combine to interpret language—including lexical and syntactic information, world knowledge, and information about others’ mental states (Garnsey et al., 1997; MacDonald et al., 1994; Tanenhaus et al., 1995; Trueswell et al., 1993, 1994)—can be used to make informed guesses about how the linguistic signal may unfold, and much behavioral and neural evidence now suggests that readers and listeners indeed engage in such predictive behavior (Altmann & Kamide, 1999; Frank & Bod, 2011; Kuperberg & Jaeger, 2016; Shain et al., 2020; Smith & Levy, 2013). An intriguing possibility is therefore that both the human language system and successful ANN models of language are optimized to predict upcoming words in the service of efficient meaning extraction.

Going beyond the broad *idea* of prediction in language, the work presented here validates, refines, and computationally implements an explicit account of predictive processing: for the first time in the neuroscience of language, we were able to accurately predict (relative to the noise ceiling) activity across voxels as well as neuronal populations in human cortex during the processing of sentences. We quantitatively test the predictive processing hypothesis at the level of voxel/electrode/fROI responses and, through the use of end-to-end models, related neural mechanisms to performance of models on computational tasks. Moreover, we were able to reject multiple alternative hypotheses about the objective of the language system: model performance on diverse benchmarks from the GLUE suite of benchmarks (A. Wang, Singh, et al., 2019), including judgments about syntactic and semantic properties of sentences, was not predictive of brain or behavioral scores. The best-performing computational models identified in this work serve as computational explanations for the entire language processing pipeline from word inputs to neural mechanisms to behavioral outputs. These best-performing models can now be further dissected, as well as tested on new diverse, linguistic inputs in future experiments, as discussed below.

### Importance of architecture

We also found that architecture is an important contributor to the models’ match to human brain data: untrained models with a trained linear readout performed well above chance in predicting neural activity, and this finding held under a series of controls to alleviate concerns that it could be an artifact of our training or testing methodologies (Fig. S9). This result is consistent with findings in models of early (Cadena et al., 2019; Cichy et al., 2016; Geiger et al., 2020) and high-level visual processing (Geiger et al., 2020) and speech perception (Millet & King, 2021), as well as recent results in natural language processing (Merchant et al., 2020; Tenney et al., 2019; Zhang & Bowman, 2018), but it raises important questions of interpretation in the context of human language. If we construe model training as analogous to learning in human development, then human cortex might already provide a sufficiently rich structure that allows for the relatively rapid acquisition of language (Carey & Bartlett, 1978; Dickinson, 1984; Heibeck & Markman, 1987). In that analogy, the human research community’s development of new architectures such as the transformer networks that perform well in both NLP tasks and neural language modeling could be akin to recapitulating evolution (Hasson et al., 2020), or perhaps, more accurately, selective breeding with genetic modification: structural changes are tested and the best-performing ones are incorporated into the next generation of models. Importantly, this process still optimizes for language modeling, only implicitly and on a different timescale from biological and cultural evolutionary mechanisms conventionally studied in brain and language.

More explicitly, but speculatively, it is possible that transformer networks can work as brain models of language even without extensive training because the hierarchies of local spatial filtering and pooling as found in convolutional as well as attention-based networks are a generally applicable brain-like mechanism to extract abstract features from natural signals. Regardless of the exact filter weights, transformer architectures build on word embeddings that capture both semantic and syntactic features of words, and integrate contextually weighted predictions across scales such that contextual dependencies are captured at different scales in different kernels. The representations in such randomized architectures could thus reflect a kind of multi-scale, spatially smoothed average (over consecutive inputs) of word embeddings, which might capture the statistical gist-like processing of language observed in both behavioral studies (Ferreira et al., 2002; Gibson et al., 2013; Levy, 2008b) and human neuroimaging (Mollica et al., 2020). The weight sharing within architectural sub-layers (“multi-head attention”) introduced by combinations of query-key-value pairs in transformers might provide additional consistency and coverage of representations. Relatedly, an idea during early work on perceptrons was to have random projections of input data into high-dimensional spaces and to then only train thin readouts on top of these projections. This was motivated by Cover’s theorem which states that non-linearly separable data can likely be linearly separated after projection into a high-dimensional space (Cover, 1965). These ideas have successfully been applied to kernel machines (Rahimi & Recht, 2009) and are more recently explored again with deep neural networks (Frankle et al., 2019); in short, it is possible that even random features with the right multiscale structure in time and space could be more powerful for representing human language than is currently understood. Finally, it is worth noting that the initial weights in the networks we study stem from weight initializer distributions that were chosen to provide solid starting points for contemporary architectures and lead to reasonable initial representations that model training further refines. These initial representations could thus include some important aspects of language structure already. A concrete test for these ideas would be the following: construct model variants that average over word embeddings at different scales and compare these models’ representations with those of different layers in untrained transformer architectures as well as the neural datasets. More detailed analyses, including minimal-pair model variant comparisons, will be needed to fully separate the representational contributions of architecture and training.

### Limitations and future directions

These discoveries pave the way for many exciting future directions. The most brain-like language models can now be investigated in richer detail, ideally leading to intuitive theories of their inner workings. Such research is much easier to perform on models than on biological systems given that all their structure and weights are easily accessible and manipulable (Cheney et al., 2017; Lindsey et al., 2019). For example, controlled comparisons of architectural variants and training objectives could define the necessary and sufficient conditions for human-like language processing (Samek et al., 2017), synergizing with parallel ongoing efforts in NLP to probe ANNs’ linguistic representations (Hewitt & Manning, 2019; Linzen et al., 2016; Tenney et al., 2020). Here, we worked with off-the-shelf models, and compared their match to neural data based on their performance on the next-word-prediction task vs. other tasks. Re-training many models on many tasks from scratch might determine which features are most important for brain predictivity, but is currently prohibitively expensive due to the vast space of hyper-parameters. Further, the fact that language modeling is inherently built into the evolution of language models by the NLP community, as noted above, may make it impossible to fully eliminate its influences on the architecture even for models trained from scratch on other tasks. Similarly, here, we leveraged existing neural datasets. This work can be expanded in many new directions, including a) assembling a wider range of publicly available language datasets for model testing (cf. vision (Schrimpf et al., 2018, 2020)); b) collecting data on new language stimuli for which different models make maximally different predictions (cf. vision; (Golan et al., 2019)), including sampling a wider range of language stimuli (e.g., naturalistic dialogs/conversations); c) modeling the fine-grained temporal trajectories of neural responses to language in data with high temporal resolution (which requires computational accounts that make predictions about representational dynamics); and d) querying models on the sentence stimuli that elicit the strongest responses in the language network to generate hypotheses about the critical response-driving feature/feature spaces, and perhaps to discover new organizing principles of the language system (cf. vision; (Bashivan et al., 2019; Ponce et al., 2019)).

One of the major limiting factors in modeling the brain’s language network is the availability of adequate recordings. Although an increasing number of language fMRI, MEG, EEG, and intracranial datasets are becoming publicly available, they often lack key properties for testing computational language models. In particular, what is needed are data with high signal-to-noise ratio, where neural responses to a particular stimulus (e.g., sentence) can be reliably estimated. However, most past language neuroscience research has focused on coarse distinctions (e.g., sentences with vs. without semantic violations, or sentences with different syntactic structures); as a result, any single sentence is generally only presented once, and neural responses are averaged across all the sentences within a ‘condition’ (in contrast, monkey physiology studies of vision typically present each stimulus dozens of times to each animal; e.g., Majaj et al., 2015). (Studies that use ‘naturalistic’ language stimuli like stories or movies also typically present the stimuli once, although naturally occurring repetitions of words / n-grams can be useful.) One of the neural datasets in the current study (Pereira2018) presented each sentence thrice to each subject and exhibited the highest ceiling (0.32; cf. Fedorenko2016: 0.17, Blank2014: 0.20). But even this ceiling is low relative to single cell recordings in the primate ventral stream (e.g., 0.82 for IT recordings; Schrimpf et al., 2018). Such high reliability may not be attainable for higher-level cognitive domains like language, where processing is unlikely to be strictly bottom-up/stimulus-driven. However, this is an empirical question that past work has not attempted to answer and that will be important in the future for building models that can accurately capture the neural mechanisms of language.

How can we develop models that are even more brain-like? Despite impressive performance on the datasets and metrics here, ANN language models are far from human-level performance in the hardest problem of language understanding. An important open direction is to integrate language models like those used here with models and data resources that attempt to capture aspects of meaning important for commonsense world knowledge (e.g., Bisk et al., 2020; Bosselut et al., 2020; Sap et al., 2019, 2020; Yi et al., 2018). Such models might capture not only predictive processing in the brain—what word is likely to come next—but also semantic parsing, mapping language into conceptual representations that support grounded language understanding and reasoning (Bisk et al., 2020). The fact that language models lack meaning and focus on local linguistic coherence (Mahowald et al., 2020; Wilcox et al., 2020) may explain why their representations fall short of ceiling on *Blank2014*, which uses story materials and may therefore require long-range contexts.

Another key missing piece in the mechanistic modeling of human language processing is a more detailed mapping from model components onto brain anatomy. In particular, aside from the general targeting of the fronto-temporal language network, it is unclear which parts of a model map onto which components of the brain’s language processing mechanisms. In models of vision, for instance, attempts are made to map ANN layers and neurons onto cortical regions (Kubilius et al., 2019) and sub-regions (Lee & DiCarlo, 2018). However, whereas function and its mapping onto anatomy is at least coarsely defined in the case of vision (Felleman & Van Essen, 1991), a similar mapping is not yet established in language beyond the broad distinction between perceptual processing and higher-level linguistic interpretation (e.g. Fedorenko & Thompson-Schill, 2014). The ANN models of human language processing identified in this work might also serve to uncover these kinds of anatomical distinctions for the brain’s language network – perhaps, akin to vision, groups of layers relate to different cortical regions and uncovering increased similarity to neural activity of one group over others could help establish a cortical hierarchy. The brain network that supports higher-level linguistic interpretation—which we focus on here—is extensive and plausibly contains meaningful functional dissociations, but how the network is precisely subdivided and what respective roles its different components play remains debated. Uncovering the internal structure of the human language network, for which intracranial recording approaches with high spatial and temporal resolution may prove critical (Mukamel & Fried, 2012; Parvizi & Kastner, 2018), would allow us to guide and constrain models of tissue-mapped mechanistic language processing. More precise brain-to-model mappings would also allow us to test the effects of perturbations on models and compare them against perturbation effects in humans, as assessed with lesion studies or reversible stimulation. More broadly, anatomically and functionally precise models are a required software component of any form of brain-machine-interface.

### Conclusions

Taken together, our findings suggest that predictive artificial neural networks serve as viable hypotheses for how predictive language processing is implemented in human neural tissue. They lay a critical foundation for a promising research program synergizing high-performing mechanistic models of natural language processing with large-scale neural and behavioral measurements of human language comprehension in a virtuous cycle of integrative modeling: testing model ability to predict neural and behavioral measurements, dissecting the best-performing models to understand which components are critical for high brain predictivity, developing better models leveraging this knowledge, and collecting new data to challenge and constrain the future generations of neurally plausible models of language processing.

## Methods

### 1. Neural dataset 1: fMRI (*Pereira2018*)

We used the data from Pereira et al.’s (2018) Experiments 2 (n=9) and 3 (n=6) (10 unique participants). (The set of participants is not identical to Pereira et al., 2018: i) one participant (tested at Princeton) was excluded from both experiments here to keep the fMRI scanner the same across participants; and ii) two participants who were excluded from Experiment 2 in Pereira et al., 2018, based on the decoding results in Experiment 1 of that study were included here, to err on the conservative side.) Stimuli for Experiment 2 consisted of 384 sentences (96 text passages, four sentences each), and stimuli for Experiment 3 consisted of 243 sentences (72 text passages, 3 or 4 sentences each). The two sets of materials were constructed independently, and each spanned a broad range of content areas. Sentences were 7-18 words long in Experiment 2, and 5-20 words long in Experiment 3. The sentences were presented on the screen one at a time for 4s (followed by 4s of fixation, with additional 4s of fixation at the end of each passage), and each participant read each sentence three times, across independent scanning sessions (see Pereira et al., 2018 for details of experimental procedure and data acquisition).

#### Preprocessing and response estimation

Data preprocessing was carried out with SPM5 (using default parameters, unless specified otherwise) and supporting, custom MATLAB scripts. (Note that SPM was only used for preprocessing and basic modeling—aspects that have not changed much in later versions; for several datasets, we have directly compared the outputs of data preprocessed and modeled in SPM5 vs. SPM12, and the outputs were nearly identical.) Preprocessing included motion correction (realignment to the mean image of the first functional run using 2nd-degree b-spline interpolation), normalization (estimated for the mean image using trilinear interpolation), resampling into 2mm isotropic voxels, smoothing with a 4mm FWHM Gaussian filter and high-pass filtering at 200s. A standard mass univariate analysis was performed in SPM5 whereby a general linear model (GLM) estimated the response to each sentence in each run. These effects were modeled with a boxcar function convolved with the canonical Hemodynamic Response Function (HRF). The model also included first-order temporal derivatives of these effects (which were not used in the analyses), as well as nuisance regressors representing entire experimental runs and offline-estimated motion parameters.

#### Functional localization

Data analyses were performed on fMRI BOLD signals extracted from the bilateral fronto-temporal language network. This network was defined functionally in each participant using a well-validated language localizer task (Fedorenko et al., 2010), where participants read sentences vs. lists of nonwords. This contrast targets brain areas that support ‘high-level’ linguistic processing, past the perceptual (auditory/visual) analysis. Brain regions that this localizer identifies are robust to modality of presentation (e.g., Fedorenko et al., 2010; Scott et al., 2017), as well as materials and task (Diachek et al., 2020). Further, these regions have been shown to exhibit strong sensitivity to both lexico-semantic processing (understanding individual word meanings) and combinatorial, syntactic/semantic processing (putting words together into phrases and sentences) (Bautista & Wilson, 2016; I. Blank et al., 2016; I. A. Blank & Fedorenko, 2020; Fedorenko et al., 2010, 2012, 2016, 2020). Following prior work, we used group-constrained, participant-specific functional localization (Fedorenko et al., 2010). Namely, individual activation maps for the target contrast (here, sentences>nonwords) were combined with “constraints” in the form of spatial ‘masks’—corresponding to data-driven, large areas within which most participants in a large, independent sample show activation for the same contrast. The masks (available from https://evlab.mit.edu/funcloc/ and used in many prior studies e.g., Jouravlev et al., 2019; Diachek et al., 2020; Shain et al., 2020) included six regions in each hemisphere: three in the frontal cortex (two in the inferior frontal gyrus, including its orbital portion: IFGorb, IFG; and one in the middle frontal gryus: MFG), two in the anterior and posterior temporal cortex (AntTemp and PostTemp), and one in the angular gyrus (AngG). Within each mask, we selected 10% of most localizer-responsive voxels (voxels with the highest *t*-value for the localizer contrast) following the standard approach in prior work. This approach allows to pool data from the same functional regions across participants even when these regions do not align well spatially. Functional localization has been shown to be more sensitive and to have higher functional resolution (Nieto-Castanon & Fedorenko, 2012) than the traditional group-averaging approach (Holmes & Friston, 1998), which assumes voxel-wise correspondence across participants. This is to be expected given the well-established inter-individual differences in the mapping of function to anatomy, especially pronounced in the association cortex (e.g., Frost & Goebel, 2012; Tahmasebi et al., 2012; Vazquez-Rodriguez et al., 2019).

We constructed a stimulus-response matrix for each of the two experiments by i) averaging the BOLD responses to each sentence in each experiment across the three repetitions, resulting in 1 data point per sentence per language-responsive voxel of each participant, selected as described above (13,553 voxels total across the 10 participants; 1,355 average, ±6 std. dev.), and ii) concatenating all sentences (384 in Experiment 2 and 243 in Experiment 3), yielding a 384×12,195 matrix for Experiment 2, and a 243×8,121 matrix for Experiment 3.

To examine differences in neural predictivity between the language network and other parts of the brain, we additionally extracted fMRI BOLD signals from two other networks: the multiple demand (MD) network (Duncan, 2010; Fedorenko et al., 2013) and the default mode network (DMN) (Buckner et al., 2008; Buckner & DiNicola, 2019). These networks were also defined functionally using well-validated localizer contrasts (Fedorenko et al., 2013; Mineroff et al., 2018) using a similar procedure as the one used for defining the language network: combining a set of ‘masks’ with individual activation maps, and selecting top 10% of most localizer-responsive voxels within each mask. Both networks were defined using a spatial working memory task (Fedorenko et al., 2011, 2013). For the MD network, we used the hard>easy contrast, and for the DMN network, we used the fixation>hard contrast. As for the language network, the MD and DMN masks were derived from large sets of participants for those contrasts, and are also available at https://evlab.mit.edu/funcloc/. The MD network and the DMN included 29,936 (2,994±230) and 10,978 (1,098±7) voxels, respectively.

### 2. Neural dataset 2: ECoG (*Fedorenko2016*)

We used the data from Fedorenko et al.’s (2016) study (n=5). (The set of participants includes one participant, S2, who was excluded from the main analyses in Fedorenko et al., 2016 due to a small number of electrodes of interest; because we here used only language-responsiveness as the criterion for electrode selection, this participant had enough electrodes to be included.) Stimuli consisted of 80 hand-constructed 8-word long semantically and syntactically diverse sentences and 80 lists of nonwords (as well as some other stimuli not used in the current study). For the critical analyses, we selected a set of 52 sentences that were presented to all participants. The materials were presented visually one word at a time (for 450 or 700 ms), and participants performed a memory probe task after each stimulus (see Fedorenko et al., 2016 for details of the experimental procedure and data acquisition).

#### Preprocessing and response estimation

We here provide only a brief summary, highlighting points of deviation from Fedorenko et al. (2016). The total numbers of implanted electrodes were 120, 128, 112, 134, and 98 for the five participants, respectively. Signals were digitized at 1200 Hz. Similar to Fedorenko et al. (2016), i) the recordings were high-pass filtered with a cut off frequency of 0.5 Hz; ii) reference, ground, and electrodes with high noise levels were removed, leaving 117, 118, 92, 130, and 88 electrodes (for these analyses, we were more permissive with respect to noise levels compared to Fedorenko et al., 2016, to include as many electrodes in the analyses as possible; hence the numbers of analyzed electrodes are higher here than in the original study for 4 of the 5 participants); iii) spatially distributed noise common to all electrodes was removed using a common average reference spatial filter between electrodes with line noise smaller than a predefined threshold (electrodes connected to the same amplifier); and iv) a set of notch filters were used to remove the 60 Hz line noise and its harmonics. To extract the high gamma band activity—which has been shown to correspond to spiking neural activity in the vicinity of the electrodes (Buzsáki et al., 2012)—we used a gaussian filter bank with centers at 73, 79.5, 87.8, 96.9, 107, 118.1, 130.4, and 144 Hz, and standard deviations of 4.68, 4.92, 5.17, 5.43, 5.7, 5.99, 6.3, and 6.62 Hz, respectively. This approach differs from Fedorenko et al. (2016), where an IIR band-pass filter was used to select frequencies in the range of 70-170 Hz, and is likely more sensitive (Dichter et al. 2018). Finally, as in Fedorenko et al. (2016), the Hilbert transform was used to extract the analytic signal (Lawrence Marple, 1999) (except here, the average of the Hilbert signal across the eight filters was used as high-gamma signal), z-scored for each electrode with respect to the activity throughout the experiment, and the signal envelopes were downsampled to 300 Hz for further analysis (we did not additionally low-pass filter at 100 Hz, as in Fedorenko et al., 2016).

#### Functional localization

Mirroring the fMRI approach, where we focused on language-responsive voxels, data analyses were performed on signals extracted from language-responsive electrodes. These electrodes were defined in each participant using the same localizer contrast as in the fMRI datasets. In particular, we examined electrodes in which the envelope of the high gamma signal was significantly higher (at p<.01) for trials of the sentence condition than the nonword-list condition (for details, see Fedorenko et al., 2016).

We constructed a stimulus-response matrix by i) averaging the *z*-scored high-gamma signal over the full presentation window of each word in each sentence, resulting in 8 data points per sentence per language-responsive electrode (97 electrodes total across the 5 participants; 47, 8, 9, 15, and 18 for participants S1 through S5, respectively), and ii) concatenating all words in all sentences (416 words across the 52 sentences), yielding a 416×97 matrix.

To examine differences in neural predictivity between language-responsive and other electrodes, we additionally extracted high gamma signals from a set of ‘stimulus-responsive’ electrodes. Stimulus-responsive electrodes were defined as electrodes in which the envelope of the high gamma signal for the sentence condition was significantly different (at p<0.05 by a paired-samples *t*-test) from the activity during the inter-trial fixation interval preceding the trial. This selection procedure resulted in 67, 35, 20, 29, and 26 electrodes. As expected, this set of electrodes included many of the language-responsive electrodes; for the analysis in SI-4, we exclude the language-responsive electrodes leaving 105 stimulus-(but not language-) responsive electrodes.

### 3. Neural dataset 3: fMRI (*Blank2014*)

We used the data from Blank et al. (2014) (n=5). (The set of participants includes 5 of the 10 participants in Blank et al., 2014, because we wanted each participant to have been exposed to the same materials and as many stories as possible; the 5 participants included here all heard eight stories.) Stimuli consisted of stories from the publicly available Natural Stories Corpus (Futrell et al., 2018). These stories, adapted from existing texts (fairy tales and short stories) were designed to be “deceptively naturalistic”: they contained an over-representation of rare words and syntactic constructions embedded in otherwise natural linguistic context. The stories were presented auditorily (each was ∼5 min in duration), and following each story, participants answered 6 comprehension questions (see Blank et al., 2014 for details of the experimental procedure, data acquisition, and preprocessing).

#### Functional localization

As in the Pereira2018 dataset, data analyses were performed on fMRI BOLD signals extracted from the language network. From each language-responsive voxel of each participant, the BOLD time-series for each story was extracted. Across the eight stories, the BOLD time-series included 1,317 time-points (TRs, time of repetition; TR=2s and corresponds to the time it takes to acquire the full set of slices through the brain). To align the neuroimaging data with the story text, we first split the text into consecutive 2-second intervals (corresponding to the fMRI TRs) based on the auditory recording; if a word straddled boundaries of intervals, it was assigned to the 2s interval in which that spoken word ended. Each of the resulting intervals thus included a story “fragment”, which could be a full short sentence, part of a longer sentence, or a transition between the end of one sentence and the beginning of another. Due to the temporal resolution of the HRF, whose peak’s latency is 4-6 seconds, we assumed that each time-point in the BOLD signal represented activity elicited by the text fragment that occurred 4s (i.e., 2 TRs) earlier.

We constructed a stimulus-response matrix by i) averaging the BOLD signals corresponding to each TR in each story across the voxels within each ROI of each participant (averaging across the voxels within ROIs was done to increase the signal-to-noise ratio), resulting in 1 data point per TR per language-responsive ROI of each participant (60 ROIs total across the 5 participants), and ii) concatenating all story fragments (1,317 ‘stimuli’), yielding a 1,317×60 matrix.

### 4. Behavioral dataset: Self-paced reading (*Futrell2018*)

We used the data from Futrell et al. (2018) (n=179). (The set of participants excludes 1 participant for whom data exclusions—see below—left only 6 data points or fewer.) Stimuli consisted of ten stories from the Natural Stories Corpus (same materials as those used in *Blank2014*, plus two additional stories), and any given participant read between 5 and all 10 stories. The stories were presented online (on Amazon’s Mechanical Turk platform) visually in a dashed moving window display—a standard approach in behavioral psycholinguistic research (Just et al., 1982). In this approach, participants press a button to reveal each consecutive word of the sentence or story; as they press the button again, the word they just saw gets converted to dashes again, and the next word is uncovered. The time between button presses provides an estimate of overall language comprehension difficulty, and has been shown to be robustly sensitive to both lexical and syntactic features of the stimuli (Grodner & Gibson, 2005; Smith & Levy, 2013, inter alia) (see Futrell et al., 2018 for details of the experimental procedure and data acquisition.) We followed data exclusion criteria in Futrell et al. (2018): for any given participant, we only included data for stories where they answered 5 or all 6 comprehension questions correctly, and we excluded reading times (RTs) that were shorter than 100 ms or longer than 3000 ms.

We constructed a stimulus-response matrix by i) obtaining the RTs for each word in each story for each participant (848,762 RTs total across the 179 participants; 338 average, ±173 std. dev.), and ii) concatenating all words in all sentences (10,256 words across 485 sentences), yielding a 10,256×179 matrix.

### 5. Computational models

We tested 43 language models that were selected to sample a broad range of computational designs across three major types of architecture: embeddings, recurrent architectures, and attention-based ‘transformer’ architectures. Here we provide a brief overview (see Table SI-10 for a summary of key features varying across the models). **GloVe** (Pennington et al., 2014) is a word embedding model where embeddings are positioned based on co-occurrence in the Common Crawl corpus; **ETM** (Dieng et al., 2019, 20ng dataset) combines word embeddings with an embedding of each word’s assigned topic; and **word2vec** (Mikolov et al., 2013)—abbreviated as w2v—provides embeddings which are trained to guess a word based on its context. **lm_1b** (Jozefowicz et al., 2016) is a 2-layer long short-term memory (LSTM) model trained to predict the next word in the One Billion Word Benchmark (Chelba et al., 2014); and the **skip-thoughts** model (Kiros et al., 2015) is trained to reconstruct surrounding sentences in a passage. For all 38 transformer models (pretrained models from the HuggingFace library (Wolf et al., 2019)), we only evaluate the encoder and not the decoder; the encoders process long contexts (100s of words) with a deep neural network stack of multiple attention heads that operate in a feed-forward manner (except the Transformer-XL-wt103 and the two XLNet models, which use recurrent processing), and differ mostly in the choice of directionality, network architecture, and training corpora (Table SI-11). We highlight key features of different classes of transformer models (BERT, RoBERTa, XLM, XLM-RoBERTa, Transformer-XL-wt103, XLNet, CTRL, T5, AlBERT, and GPT) in the order in which they appear in the bar-plots (e.g., Fig. 2a), except for the three ‘distilled’ models (Sanh et al., 2019), which we mention in the end. **BERT** transformers (Devlin et al., 2018) (n=4; bert-base-uncased, bert-base-multilingual-cased, bert-large-uncased, bert-large-uncased-whole-word-masking) are optimized to train bidirectional representations taking into account context both to the left and right of a masked token. **RoBERTa** transformers (Liu et al., 2019) (n=2; roberta-base, roberta-large) as a variation of BERT improve training hyper-parameters such as masking tokens dynamically instead of always masking the same token. **XLM** models (Lample & Conneau, 2019) (n=7; xlm-mlm-enfr-1024, xlm-clm-enfr-1024, xlm-mlm-xnli15-1024, xlm-mlm-100-1280, xlm-mlm-en2048) learn cross-lingual models by predicting the next (“clm”) or a masked (“mlm”) token in a different language. **XLM-RoBERTa** (Conneau et al., 2019) (n=2; xlm-roberta-base, xlm-roberta-large) combines RoBERTa masking with cross-lingual training in XLM. **Transformer-XL-wt103** (Dai et al., 2020) adds a recurrence mechanism to GPT (see below) and trains on the smaller WikiText-103 corpus. **XLNet** transformers (Yang et al., 2019) (n=2; xlnet-base-cased, xlnet-large-cased) permute tokens in a sentence to predict the next token. **CTRL** (Keskar et al., 2019) adds control codes to GPT (see below) which influence text generation in a specific style. **T5** transformers (Raffel et al., 2019) (n=5; t5-small, t5-base, t5-large, t5-3b, t5-11b) train the same model across a range of tasks including the prediction of multiple corrupted tokens, GLUE (A. Wang, Singh, et al., 2019), and SuperGLUE (A. Wang, Pruksachatkun, et al., 2019) in a text-to-text manner where the task is provided as a text prefix. **AlBERT** transformers (Lan et al., 2019) (n=8; albert-base-v1, albert-large-v1, albert-xlarge-v1, albert-xxlarge-v1, albert-base-v2, albert-large-v2, albert-xlarge-v2, albert-xxlarge-v2) use parameter-sharing and model inter-sentence coherence. **GPT** transformers (n=5) are trained to predict the next token in a large dataset emphasizing document quality (openaigpt (Radford et al., 2018) on the Book Corpus dataset, gpt2, gpt2-medium, gpt2-large, and gpt2-xl (Radford et al., 2019) on WebText). Finally, **distilled versions** of models (Sanh et al., 2019) (n=3; distilbert-base-uncased, distilgpt2, distilroberta-base) train compressed models on a larger teacher network.

To retrieve model representations, we treated each model as an experimental participant (Figure 1) and ran the same experiment on it that was run on humans. Specifically, sentences were fed in sequentially into the model (for Pereira2018, Blank2014, and Futrell2018, sentences were grouped by topic / story to approximate the procedure with human participants). For embedding and recurrent models, sentences were fed in word-by-word; for transformers, the context before (but not after) each word was also fed into the models due to their lack of memory; the length of the context was determined by the models’ architectures. For recurrent models, the memory was reset after each story (*Pereira2018, Blank2014* and *Futrell2018*), or each sentence (*Fedorenko2016*).

After the processing of each word, we retrieved (“recorded”) model representations at every computational block (e.g., one LSTM cell or one Transformer encoder block). (Word-by-word processing increases computational cost but is necessary to avoid bidirectional models, like the BERT transformers, seeing the future.) When comparing against human recordings spanning more than one word such as a sentence (*Pereira2018*) or story fragment (*Blank2014*), we aggregated model representations: for the embedding models, we used the mean of the word representations; for recurrent and transformer models, we used the representation of the last word since these models already aggregate representations of the preceding context, up to a maximum context length of 512 tokens.

### 6. Comparison of models to brain measurements

We treated the model representation at each layer separately and tested how well it could predict human recordings (for *Pereira2018*, we treated the two experiments separately, but averaged the results across experiments for all plots except Fig. 2c). To generate predictions, we used 80% of the stimuli (sentences in *Pereira2018*, words in *Fedorenko2016* and *Futrell2018*, and story fragments in *Blank2014*; Fig. 1) to fit a linear regression from the corresponding 80% of model representations to the corresponding 80% of human recordings. We applied the regression on model representations of the held-out 20% of stimuli to generate model predictions, which we then compared against the held-out 20% of human recordings with a Pearson correlation. This process was repeated five times, leaving out different 20% of stimuli each time, and we computed the per-voxel/electrode/ROI mean predictivity across those five splits. We aggregated these per-voxel/electrode/ROI scores by taking the median of scores for each participant’s voxels/electrodes/ROIs and then computing the median and median absolute deviation (m.a.d.) across participants (over per-participant scores). Finally, this score was divided by the estimated ceiling value (see <u>Estimation of ceiling</u> below) to yield a final score in the range of [0, 1]. We report the results for the best-performing layer for each model (SI-12) but controlled for the generality of layer choices in train/test splits (Fig. S2b,c).

### 7. Estimation of ceiling

Due to intrinsic noise in biological measurements, we estimated a ceiling value to reflect how well the best possible model of an average human could perform. To do so, we first subsampled—for each dataset separately— the data with n recorded participants into all possible combinations of s participants for all s ∈ [2, *n*] (e.g. {2, 3, 4, 5} for *Fedorenko2016* with n=5 participants). For each subsample s, we then designated a random participant as the target that we attempt to predict from the remaining s − 1 participants (e.g., predict 1 subject from 1 (other) subject, 1 from 2 subjects, …, 1 from 4, to obtain a mean score for each voxel/electrode/ROI in that subsample. To extrapolate to infinitely many humans and thus to obtain the highest possible (most conservative) estimate, we fit the equation 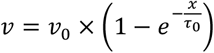 where x is each subsample’s number of participants, v is each subsample’s correlation score and *v*_o_ and *τ*_o_ are the fitted parameters for asymptote and slope respectively. This fitting was performed for each voxel/electrode/ROI independently with 100 bootstraps each to estimate the variance where each bootstrap draws x and v with replacement. The final ceiling value was the median of the per-voxel/electrode/ROI ceilings *v*_o_.

For *Fedorenko2016*, a ceiling was estimated for each electrode in each participant, so each electrode’s raw value was divided by its own ceiling value. Similarly, for *Blank2014*, a ceiling was estimated for each ROI in each participant, so each ROI’s raw value was divided by its own ceiling value. For *Pereira2018*, we treated the two experiments separately, focusing on the 5 participants that completed both experiments to obtain full overlap in the materials for each participant, and used 10 random sub-samples to keep the computational cost manageable. A ceiling was estimated for all voxels in the 5 participants who participated in both experiments. Each voxel’s raw predictivity value was divided by the average ceiling estimate (across all the voxels for which it was estimated). For *Futrell2018*, given the large number of participants and because most participants only had measurements for a subset of the stimuli, we did not hold out one participant but rather tested how well the mean RTs for one half of the participants predicted the RTs for the other half of participants. We further took 5 random subsamples at every 5 participants, starting from 1, and built 3 random split-halves, again to keep computational cost manageable. A ceiling was estimated for each participant, and each participant’s raw values were divided by this ceiling. (Note that this approach is even more conservative than the leave-one-out approach, because split-half correlations tend to be higher than one-vs.-rest, due to a reduction in noise when averaging (for each half).)

### 8. Language Modeling

To assess the models’ performance on the normative next-word-prediction task, we used a dataset of 720 Wikipedia articles, WikiText-2 (Merity et al., 2016), with 2M training, 218k validation, and 246k test tokens (words and word-parts). These tokens were processed by model-specific tokenization with a maximum vocabulary size of 250k, selected based on the tokens’ frequency in the model’s original training dataset, and split up into blocks of 32 tokens each (both the vocabulary size and the length of blocks were constrained by computational cost limitations). We sequentially fed the tokens into models as explained in <u>Methods_5</u> (<u>Computational Models)</u> and captured representations at each step from each model’s final layer (penultimate layer before the classifier if the model has a readout). To predict the next word, we fit a linear decoder from those representations to the next token over words in the vocabulary (n=50k), on the training tokens. This decoder is trained with a cross-entropy-loss 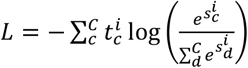 where 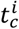 is the true label for class c and sample i, and 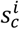 is the predicted probability of that class; the linear weights are updated with AdamW and a learning rate of 5e-5 in batches of 4 blocks until convergence as defined on the validation set. Importantly, note that we only trained weights of a readout decoder, *not* the weights of models themselves, in order to maintain the same model representations that we used in model-to-brain and model-to-behavior comparisons. The final language modeling score is reported for each model as the perplexity, i.e. the exponent of the cross-entropy loss, on the held-out test set. We ensured that our pipeline could reproduce the lower perplexity values in e.g. (Radford et al., 2019) by fine-tuning the entire model and increasing the batch size. To be able to test all models under the same conditions and with fixed representations that were used for brain prediction, we however had to use a lower batch size and only train a linear readout without fine-tuning which leads to the lower perplexity scores reported in Fig. 3. T5-11b is not part of this analysis because of lack of computational resources to run the model.

### 9. Statistical tests

As a primary metric, model-to-brain predictivity scores are reported as the Pearson correlation coefficient (denoted by “r”). These correlation scores were obtained from aggregating over individual per-voxel/electrode/ROI scores. To avoid the assumption that the neural scores are Gaussian distributed, we aggregated these per-voxel/electrode/ROI scores by taking the *median* of scores for each participant’s voxels/electrodes/ROIs and then computing the median and median absolute deviation (m.a.d.) across participants.

In addition to reporting an aggregated score across datasets, we show individual scores per dataset (visualized as bar plot insets). To obtain an error estimate for the correlation scores, we report the bootstrapped correlation coefficient, as computed by leaving out 10% of the scores and computing the r-value on the remaining 90% held-out scores (over 1,000 iterations).

All p-values less than 0.05 are summarized with one asterisk, p-values less than 0.005 with two asterisks, p-values less than 0.0005 with three asterisks, and p-values less than 0.00005 are denoted by four asterisks.

For interaction tests, we used two-sided t-tests with 1,000 bootstraps and 90% of samples per bootstrap.

## Author contributions

M.S. and J.T. conceived of the project.

M.S., I.B., G.T., C.K., N.K., J.T. and E.F. developed analyses.

I.B., G.T., M.S., C.K., E.H. and E.F. analyzed neural and behavioral data.

M.S., C.K. and G.T. implemented models.

M.S. and C.K. implemented language modeling and GLUE benchmarks.

M.S., G.T. and C. K. carried out analyses to relate model representations to neural and behavioral data.

M.S., I.B., G.T., C.K., E.H., N.K., J.T. and E.F. discussed results.

M.S., I.B., G.T., C.K., N.K., J.T. and E.F. contributed to the manuscript.

## Acknowledgments

We would like to thank Roger Levy, Steve Piantadosi, Cory Shain, Noga Zaslavsky, Antoine Bosselut, and Jacob Andreas for comments on the manuscript, Tiago Marques for comments on ceiling estimates and feature analysis, Jon Gauthier for comments on language modeling, Bruce Fischl and Ruopeng Wang for adding a Freeview functionality. MS was supported by a Takeda Fellowship, the Massachusetts Institute of Technology Shoemaker Fellowship, and the SRC Semiconductor Research Corporation. GT was supported by the MIT Media Lab Consortia and the Massachusetts Institute of Technology Singleton Fellowship. CK was funded by the Massachusetts Institute of Technology Presidential Graduate Fellowship. EH was supported by the Friends of the McGovern Institute Fellowship. NK and JT were supported by the Center for Brains, Minds, and Machines (CBMM), funded by NSF STC CCF-1231216. EF was supported by NIH awards R01-DC016607 and R01-DC016950, and by funds from the Brain and Cognitive Sciences Department and the McGovern Institute for Brain Research at MIT.

## Supplement

**S1**: Ceiling estimates for neural and behavioral datasets

**S2**: Scores generalize across metrics and layers

**S3**: Brain surface visualization of model predictivity scores

**S4**: Language specificity

**S5**: Model performance on diverse language tasks vs. model-to-brain fit

**S6**: Model’s neural predictivity for each dataset is correlated with behavioral predictivity

**S7**: Performance on next-word prediction selectively predicts model-to-behavior fit.

**S8**: Model architecture contributes to brain predictivity and untrained performance predicts trained performance

**S9**: Controls for untrained models

**S10**: Effects of model architecture and training on neural and behavioral scores

**S11**: Overview of model designs

**S12**: Distribution of layer preference (best performing layer) per voxel for GPT2-xl for *Pereira2018*

**Figure S1:**
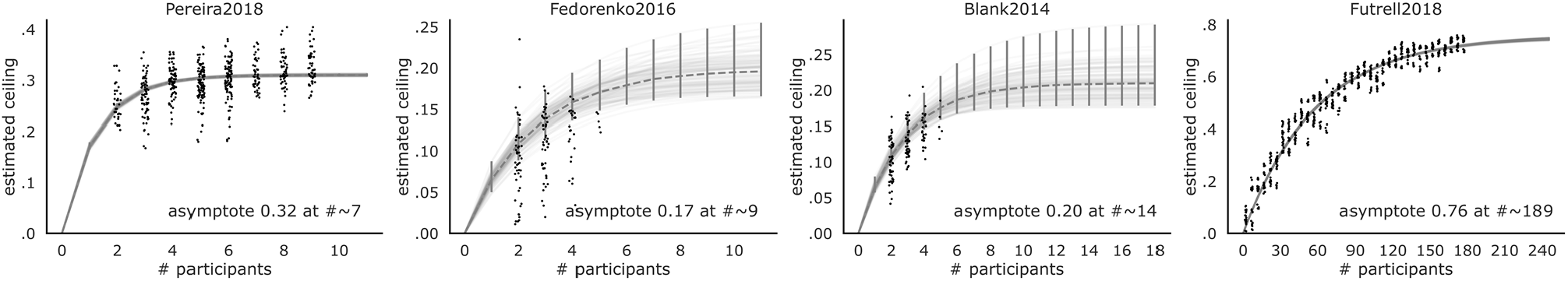
Ceiling estimates for neural and behavioral datasets. Due to intrinsic noise in biological measurements, we estimated a ceiling value to reflect how well the best possible model of an average human could perform, based on sub-samples of the total set of participants (see <u>Methods-7</u>). For each sub-sample, *s* − 1 participants are used to predict a held-out participant (except in *Futrell2018*, where this is done on split-halves, as described in the text). Each dot represents a correlation between the average scores of the *s* − 1 participants and the left-out participant for a random sub-sample of the number of participants *s* indicated on the x-axis. We then bootstrapped 100 random combinations of those dots to extrapolate (gray lines) the highest possible ceiling if we had an infinite number of participants at our disposal. The parameters of these bootstraps are then aggregated by taking the median to compute an overall estimated ceiling (dashed gray line with 95% CI in error-bars). We use this estimated ceiling to normalize model scores and here also report the number of participants at which the estimated ceiling would be met (which show that for *Pereira2018* and *Futrell2018*, the number of participants we have is at and close to the asymptote value, respectively). Ceiling levels are .32 (*Pereira2018*), .17 (*Fedorenko2016*), .20 (*Blank2014*), and .76 (*Futrell2018*).

**Figure S2:**
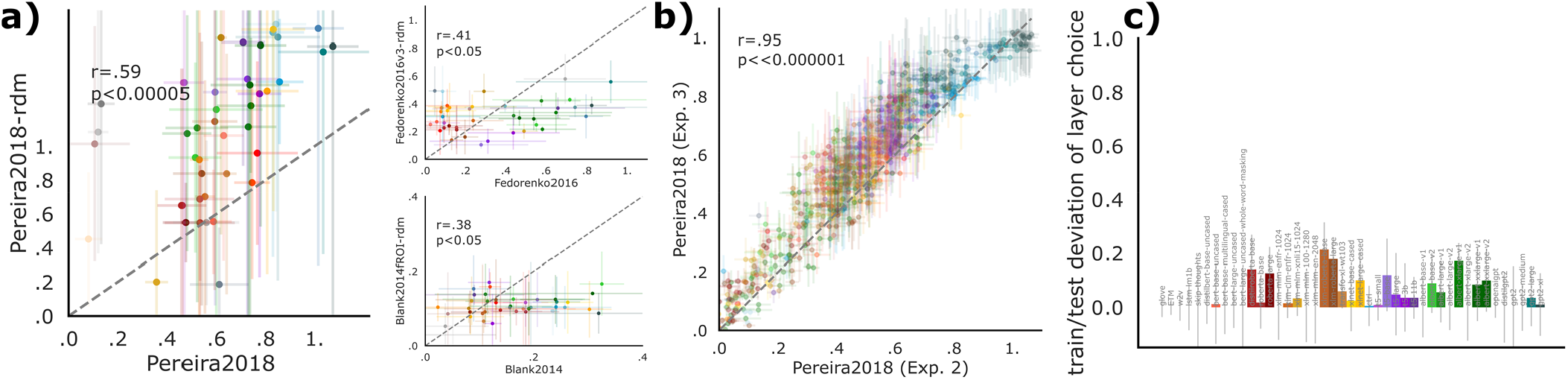
Scores generalize across metrics and layers. **a)** Model scores on each dataset generalize across different choices of a similarity metric; here we plot the predictivity metric used in the manuscript on the x-axis against a model-to-brain similarity metric based on representational dissimilarity matrices (RDMs) between models and neural representations on the y-axis. Like in the predictivity metric, stimuli along with corresponding model activations and brain recordings were split 5-fold but we then only compared the respective test splits given that the RDM metric does not employ fitting. Specifically, we followed (Kriegeskorte, 2008) and computed the RDM for each model’s activations, and a separate RDM for each brain recording dataset, based on 1 minus the Pearson correlation coefficient between pairs of stimuli; then, we measured model-brain similarity via Spearman correlation across the two RDMs’ upper triangles. The RDM score for one model on one human dataset is then the mean over splits. We ran each model and compared resulting scores with the primarily used scores from the predictivity metric. Correlations for models’ scores between the predictivity and the RDM metrics are: Pereira2018 r=.57, p<0.0001; Fedorenko2016 r=.40, p<.01; Blank2014 r=.38, p<.05. **b)** Model scores per layer generalize across dataset splits; for every layer in each model we plot its brain score (using the predictivity metric) on two experimental splits (experiment 2 and 3) of the *Pereira2018* dataset. Scores are very strongly correlated (r=.95, p<<0.000001), indicating that choosing a model’s layer on a separate dataset split will generalize to a held-out test split. **c)** Choice of layer generalizes across dataset splits; for each model we plot the difference between its score on *Pereira2018* experiment 3 when choosing the layer on experiment 3 directly (i.e. the max due to layer choice on “test set”) and its score on experiment 3 when choosing the layer on experiment 2 (choice on “train set”). The layer is chosen based on the model’s maximum score across layers on the respective dataset split. Deviations between choosing the layer on a train or test set are minimal with error bars overlapping 0, indicating that there is no substantial difference between the two choices.

**Figure S3:**
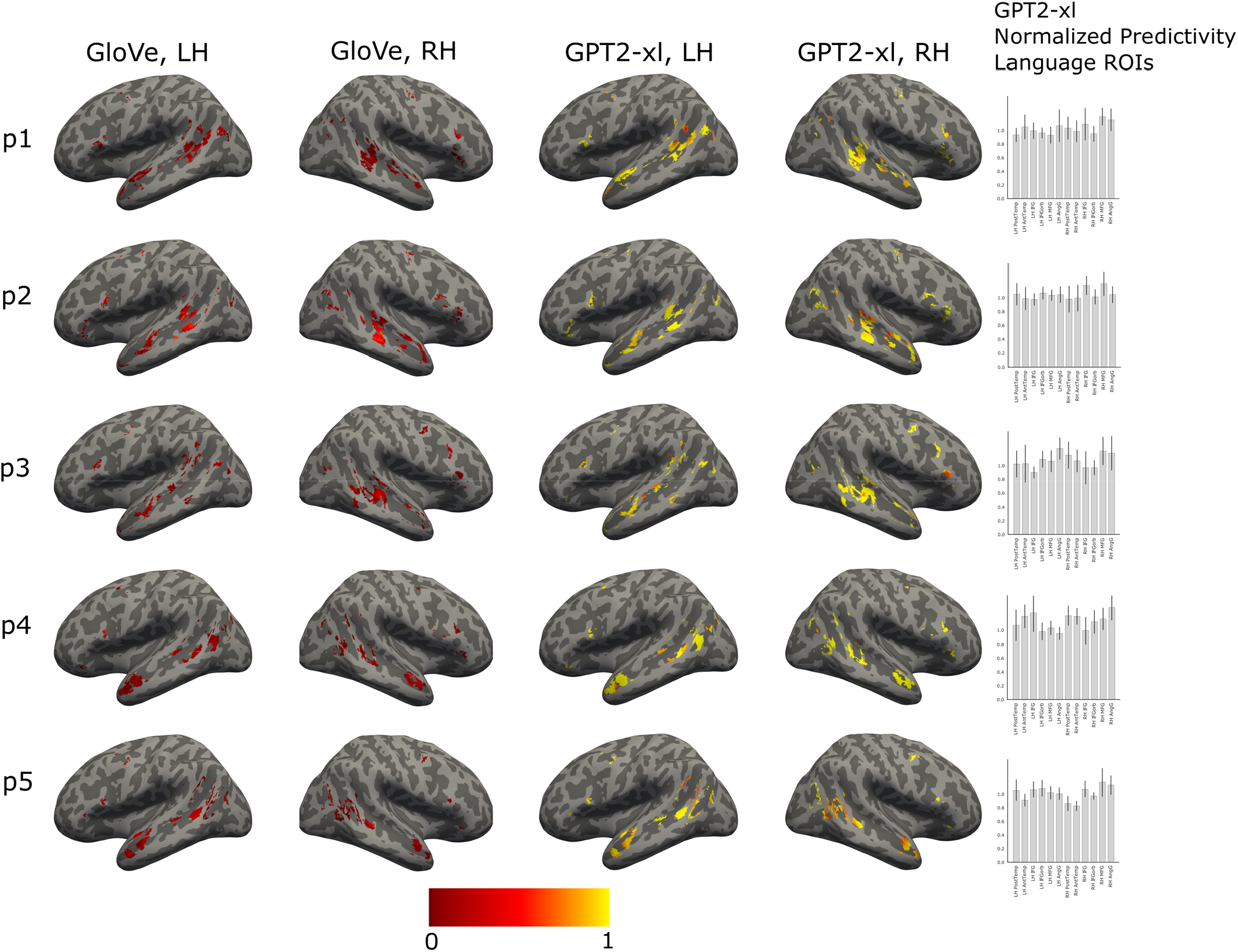
Brain surface visualization of model predictivity scores. Plots show surface projections of volumetric individual language-responsive functional ROIs in the left and right hemispheres (LH and RH) for five representative participants from *Pereira2018*. In each voxel of each fROI, we show a normalized predictivity value for two models that differ substantially in their ability to predict human data: GloVe (first two columns) and GPT2-xl (second two columns; for GPT2-xl, we show predictivity values from the overall best-performing layer, in line with how we report the results in the main text). (Note that the voxel locations are identical between GloVe and GPT2-xl, and are determined by an independent functional language localizer as described in the text; we here illustrate the differences in predictivity values, along with showing sample fROIs used in our analyses). Predictivity values were ceiling-normalized for each participant and each of 12 ROIs separately (a slight deviation from the approach in the main analysis, which was designed to control for between-region differences in reliability). The data were analyzed in the volume space and co-registered using SPM12 to Freesurfer’s standard brain CVS35 (combined volumetric and surface-based (CVS)) in the MNI152 space using nearest neighbor interpolation and no smoothing. The ceiled predictivity maps for the language localizer contrast (10% of most language-responsive voxels in each ‘mask’; <u>Methods-1</u>) were projected onto the cortical surface using mri_vol2surf in Freesurfer v6.0.0 with a projection fraction of 1. The surface projections were visualized on an inflated brain in the MNI152 space using the developer version of Freeview (assembly March 10th, 2020). The bar plots in the rightmost column show the normalized predictivity values per ROI (median across voxels) in the language network for GPT2-xl. Error bars denote m.a.d. across voxels. The distribution of predictivity values across the language-responsive voxels, and the similar predictivity magnitudes across the ROIs in the bar graphs, both suggest that the results (between-model differences in neural scores) are not driven by one particular region of the language network, but are similar across regions, and between the LH and RH components of the network (see also SI-4).

### SI-4 – Language specificity

In the analyses reported in the manuscript, we focused on the language-responsive regions / electrodes. Here, for two datasets, we investigated the model-brain relationship outside the language network in order to assess the spatial specificity of our results, i.e., to test whether they obtain only, or more strongly, in the language network compared to other parts of the brain. For both datasets, we report analyses based on *raw predictivity values*, without normalizing by the estimated noise ceiling because the brain regions of the language network differ from other parts of the brain in how strongly their activity is tied to stimulus properties during comprehension (e.g., I. A. Blank & Fedorenko, 2017, 2020; Diachek et al., 2020; Shain et al., 2020; Wehbe et al., 2020). This variability is important to take into account when comparing between functionally different brain regions/electrodes because we are interested in how well the models explain linguistic-stimulus-related neural activity. When we normalize the neural responses of a non-language-responsive region/electrode using a language comprehension task, we’re effectively isolating whatever little *stimulus-related activity* this region/electrode may exhibit, putting them on ~equal or similar footing with the language-responsive regions/electrodes. (For completeness and ease of comparison with the main analyses, we also report analyses based on normalized predictivity values.)

### Fedorenko2016

The scores obtained from language-responsive electrodes were compared to those obtained from stimulus-responsive electrodes, excluding the language-responsive ones (see <u>Methods-2</u>), for all 43 models. The number of language-responsive electrodes across five participants was 97, and the number of stimulus-, but not language-, responsive electrodes across the participants was comparable (n=105). The analysis was identical to the main analysis (see <u>Methods</u>), besides omitting the ceiling normalization for the raw predictivity analyses. As described in Methods, normalization was performed for each electrode in each participant separately.

For raw predictivity, neural responses in the language-responsive electrodes were predicted 49.21% better on average across models than the non-language-responsive electrodes (independent-samples two-tailed t-test: t=3.4, p=0.001). (For normalized predictivity, neural responses in the language-responsive electrodes were predicted 59.26% better on average across models than the non-language-responsive electrodes (t=2.24, p=0.03).)

### Pereira2018

The scores obtained from the language network were compared to those obtained from two control networks: the multiple demand (MD) network and the default mode network (DMN) (see <u>Methods</u>), for all 43 models. The number of voxels in the language network across participants was, on average, 1,355 (± 7 SD across participants), and the average number of voxels in the MD network and the DMN was comparable (MD: 2,994±230); DMN: 1,098±7). The analysis was identical to the main analysis (see <u>Methods</u>), besides omitting the ceiling normalization for the raw predictivity analyses. For the normalized predictivity analyses, the network predictivity values were normalized by their respective network ceiling values.

For raw predictivity, neural responses in the language network ROIs were predicted 16.96% better on average across models than the MD network ROIs (independent-samples two-tailed t-test: t=2.26, p=0.03) and numerically (14.33%) better than the DMN ROIs (t=1.78, p=0.08). (For normalized predictivity, neural responses in the language network ROIs were predicted numerically (6.47%) worse on average than the MD network ROIs (t=-0.92, p=0.36) and also numerically (1.05%) worse than the DMN ROIs (t=-0.31, p=0.76).)

These results suggest that—when allowing for inter-regional differences in the reliability of language-related responses—the model-to-brain relationship is stronger in the language-responsive regions/electrodes. However, we leave open the possibility that language models also explain neural responses outside the boundaries of the language network, perhaps because these models capture some parts of our general semantic knowledge, which is plausibly stored in a distributed fashion across the brain. For example, several earlier studies used simple embedding models to decode linguistic meaning from fMRI data (e.g., Wehbe et al., 2014; Huth et al., 2016; Anderson et al., 2017; Pereira et al., 2018) and reported reliable decoding not only within the language network, but also across other parts of association cortex. Given that we know that different large-scale cortical networks differ functionally in important ways (e.g., see Fedorenko & Blank, 2020, for a recent discussion of the language vs. MD networks), it will be important to investigate in future work the precise mapping between the language models’ representations and neural responses in these different functional networks.

### SI-5 – Model performance on diverse language tasks vs. model-to-brain fit

To test whether the next-word prediction task is special in predicting model-to-brain fit, we used the *Pereira2018* dataset to examine the relationship between the models’ performance on diverse language processing tasks from the General Language Understanding Evaluation (GLUE) benchmarks (Wang et al., 2018) and neural predictivity. We used a subset of the high-performing, transformer models (n=30 of the 38 where we could find published commitments of which features to use for GLUE). The GLUE benchmark encompasses nine tasks that can be classified into three categories: single-sentence judgment tasks (n=2), sentence-pair semantic similarity judgment tasks (n=3), and sentence-pair inference tasks (n=4). The two single-sentence tasks are both binary classification tasks: models are asked to determine whether a given sentence is grammatical or ungrammatical (Corpus of Linguistic Acceptability, *CoLA* (Warstadt et al., 2018)), or whether the sentiment of a sentence is positive or negative (Stanford Sentiment Treebank, *SST-2* (Socher et al., 2013)). In the semantic similarity tasks, models are asked to assert or deny the semantic equivalence of question pairs (Quora Question Pairs, *QQP* (Chen et al., 2018)) or sentence pairs (Microsoft Research Paraphrase Corpus, *MRPC* (Dolan & Brockett, 2005)), or to judge the degree of semantic similarity between two sentences on a scale of 1-5 (Semantic Textual Similarity Benchmark, *STS-B* (Cer et al., 2017)). Lastly, the benchmark contains four inference tasks, of which we include three (following Devlin et al., 2018), we exclude the Winograd Natual Language Inference, *WNLI*, task; see (12) in https://gluebenchmark.com/faq). In two of these tasks, models are asked to determine the entailment relationship between sentences in a pair using either tertiary classification: entailment, contradiction, neutral (Multi-Genre Natural Language Inference corpus, *MNLI* (Williams et al., 2018)), or binary classification: entailment or no entailment (Recognizing Textual Entailment, *RTE* (Dagan et al., 2006, Bar Haim et al., 2006, Giampiccolo et al., 2007, Bentivogli et al., 2009)). And in the third inference task, the Question Natural Language Inference, *QNLI*, task (Rajpurkar et al., 2016, White et al., 2017, Demszky et al., 2018), models are presented with question-answer pairs and asked to decide whether or not the answer-sentence contains the answer to the question.

In order to evaluate model performance on GLUE benchmark tasks, each GLUE dataset was first converted into a format that is compatible with transformer model input using functionality from the GLUE data processor provided by Huggingface transformers (https://huggingface.co/transformers/). In particular, each set of materials is represented as a matrix that includes the following dimensions: item (and sentence for multi-sentence materials) ID, ID for each individual word (with reference to the vocabulary used by the transformer models), the label (e.g., grammatical vs. ungrammatical), and the ‘attention mask’ which specifies which part(s) of the sentences the model should pay attention to (e.g., some ‘padding’ is commonly used to equalize the lengths of sentences/items to the target length of 128 tokens (again constrained by computational cost), and the attention mask is set to include only the actual words in the materials, and not the padding, and in some models to further constain which parts of the input to attend to—e.g., in GPT2 models, the rightward context is ignored). Next, each GLUE dataset was then fed into each model to obtain a sequence of hidden states at the output of the last layer of the model. Following default settings from Huggingface transformers, from these hidden states, we then extracted the token of interest: for bidirectional models such as BERT, this was the first input token—a special token ([cls]) that is appended to each item and designed for sequence classification tasks, and for unidirectional models such as GPT-2, XLNet or CTRL, this token corresponded to the last attended token (e.g., the last word/word-part in the sentence). In order to ensure a fair comparison between the models and to avoid the skewing of representations by individual task pre-training, dense linear pooling projection layers (specific to some transformer) are disregarded. Finally, we fit a linear decoder from the features of the extracted tokens of interest to the task label(s). For tasks with two or more labels, a cross-entropy loss function is used; for the task that uses a rating scale, the decoder is trained with a mean-square error (MSE) loss function. Similar to the next-word prediction task, the linear weights are updated with the AdamW optimizer and a learning rate of 5e-5 in batches of 8 blocks until convergence as defined on the validation set. Importantly, and also similar to the next-word-prediction task, we only trained weights of a readout decoder, *not* the weights of models themselves, in order to maintain the same model representations that we used in model-to-brain and model-to-behavior comparisons. To account for potential bias in the GLUE datasets, multiple metrics within tasks, as well as different metrics across tasks are reported in the GLUE benchmark. Following standards in the field, we follow GLUE evaluation metrics (A. Wang, Singh, et al., 2019) and report the final task score as accuracy for *SST-2, MNLI, RTE*, and *QNLI*, Matthew’s Correlation for *CoLA*, the average of accuracy and F1 score for MRPC, and QQP, and the average of Pearson and Spearman correlation for *STS-B*. The results are shown in Fig. S5. None of the tasks significantly predicted neural scores, suggesting that next-word prediction may be special in its ability to predict brain-like processing. As with language modeling, we were unable to evaluate T5-11b on these benchmarks due to lack of computational resources.

**Figure S5:**
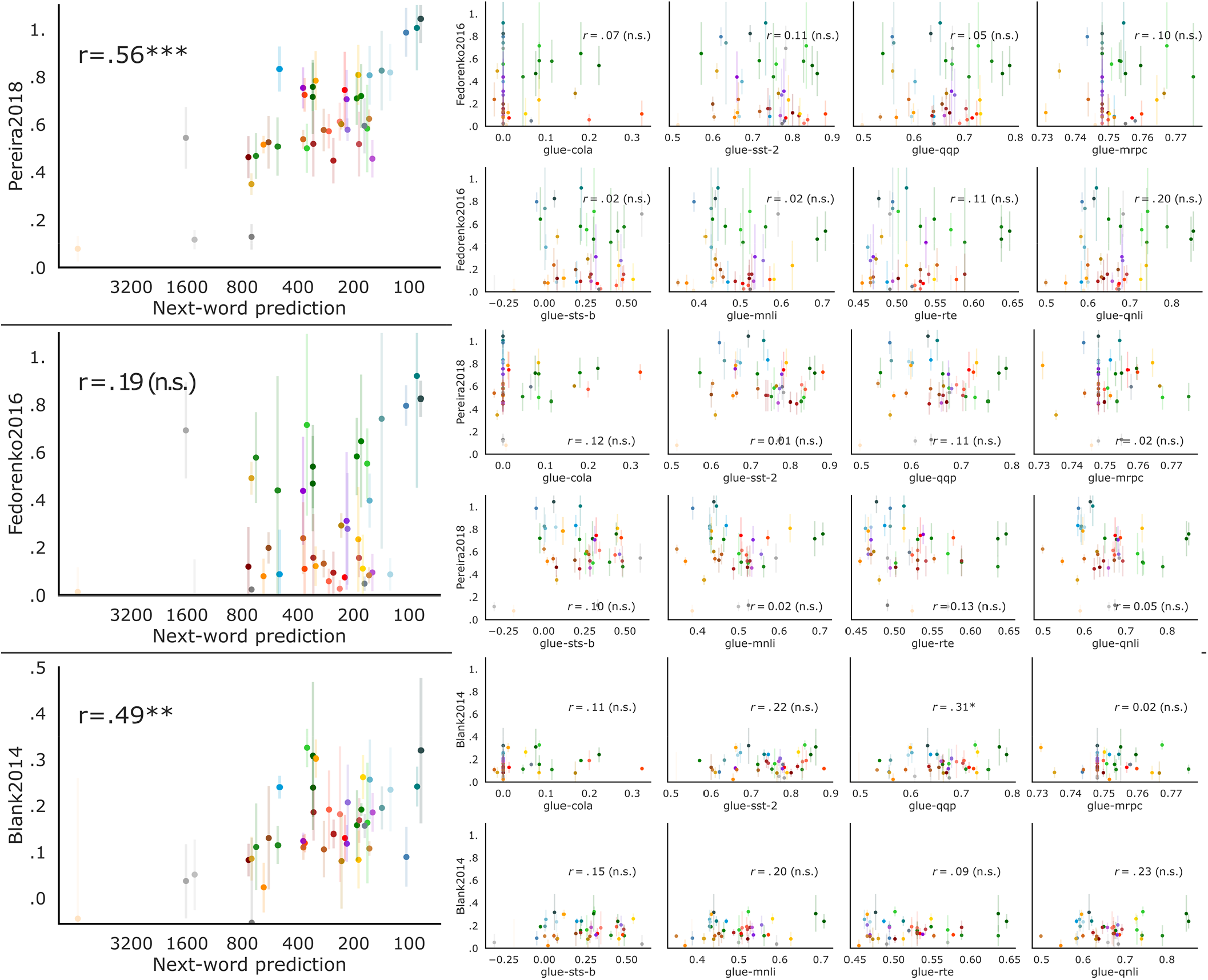
Performance on next-word prediction selectively predicts model-to-brain fit. Performance on GLUE tasks was evaluated as described in SI-5. Only the next-word prediction correlations but none of the GLUE correlations were significant.

**Figure S6:**
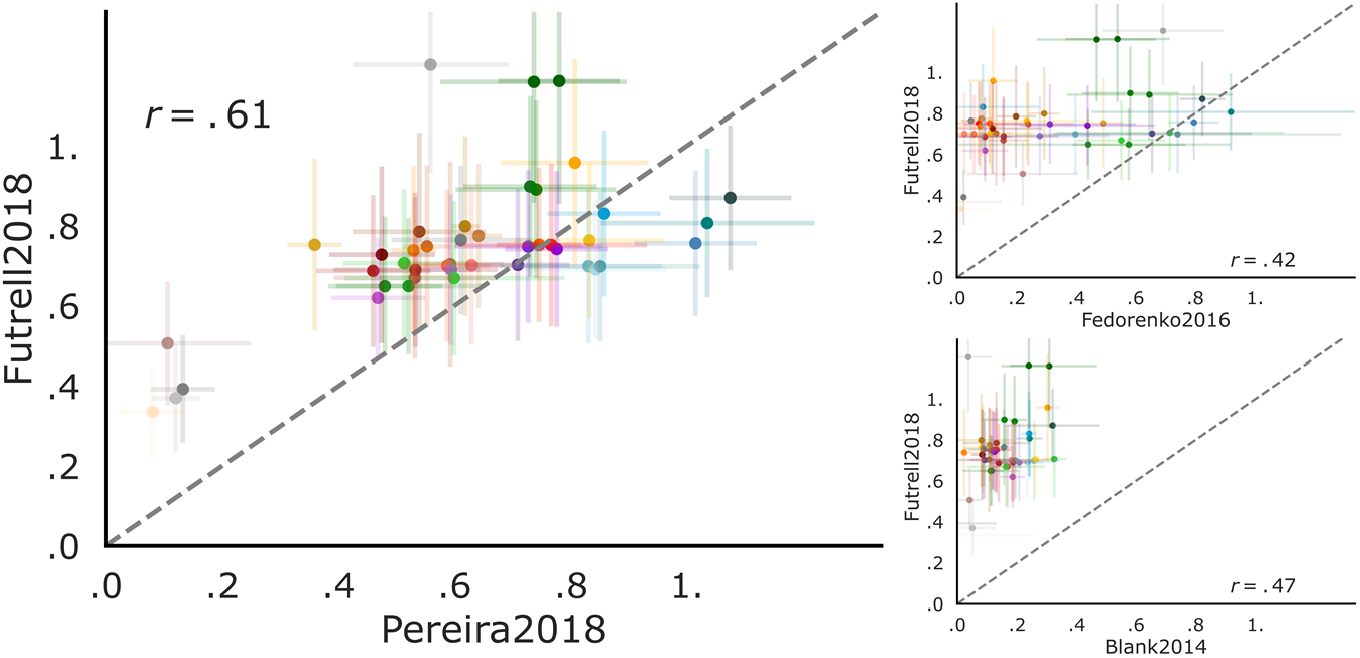
Models’ neural predictivity for each dataset is correlated with behavioral predictivity. In Fig. 4b, we showed that the models’ neural predictivity (averaged across the three neural datasets: Pereira2018, Fedorenko2016, Blank2014) correlates with behavioral predictivity. Here, we show that this relationship also holds for each neural dataset individually: Pereira2018: p<0.0001, Fedorenko2016: p<0.01, Blank2014: p<0.01.

**Figure S7:**
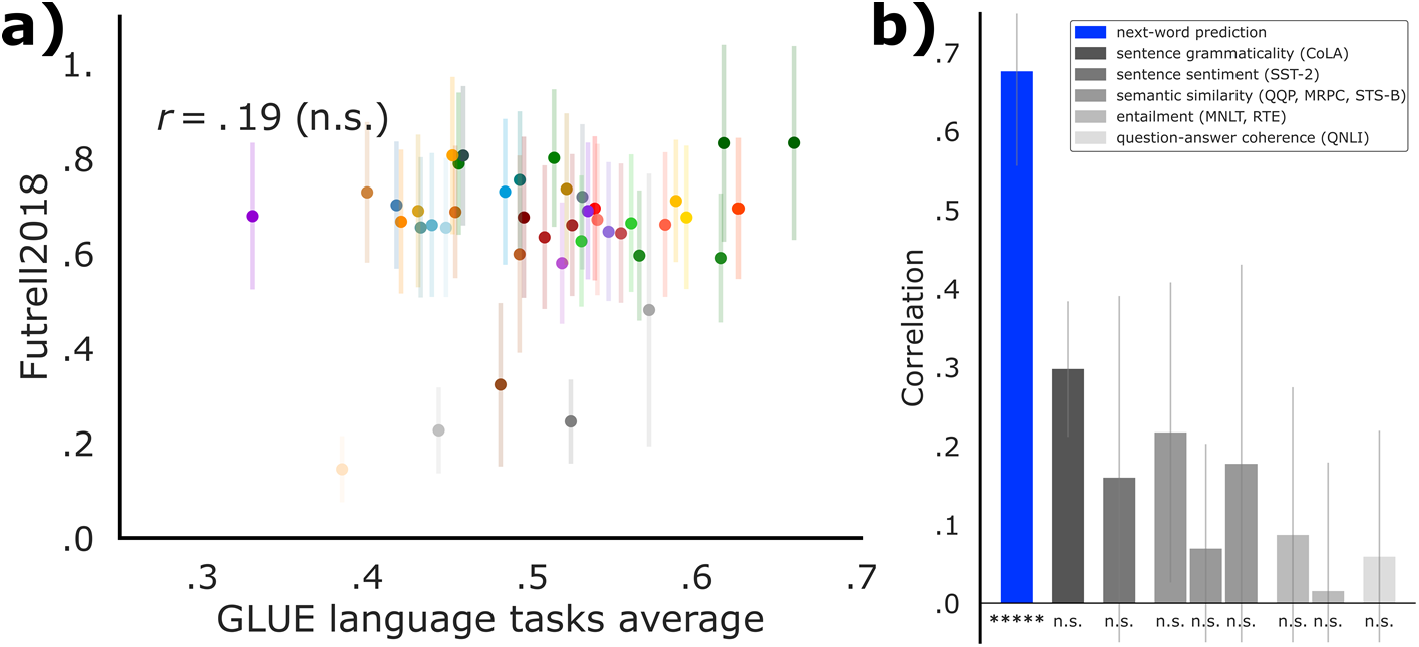
Performance on GLUE tasks does *not* predict model-to-behavior fit. In Fig. 4c, we showed a significant positive correlation of next-word prediction performance with predictivity on behavioral reading times. Here we test whether performance on GLUE tasks predicts behavioral scores (performance on GLUE tasks was evaluated as described in SI-5). Only the next-word prediction correlations but none of the GLUE correlations were significant. Notations as in Figure 3 for the GLUE average (a) and individual tasks (b).

**Figure S8:**
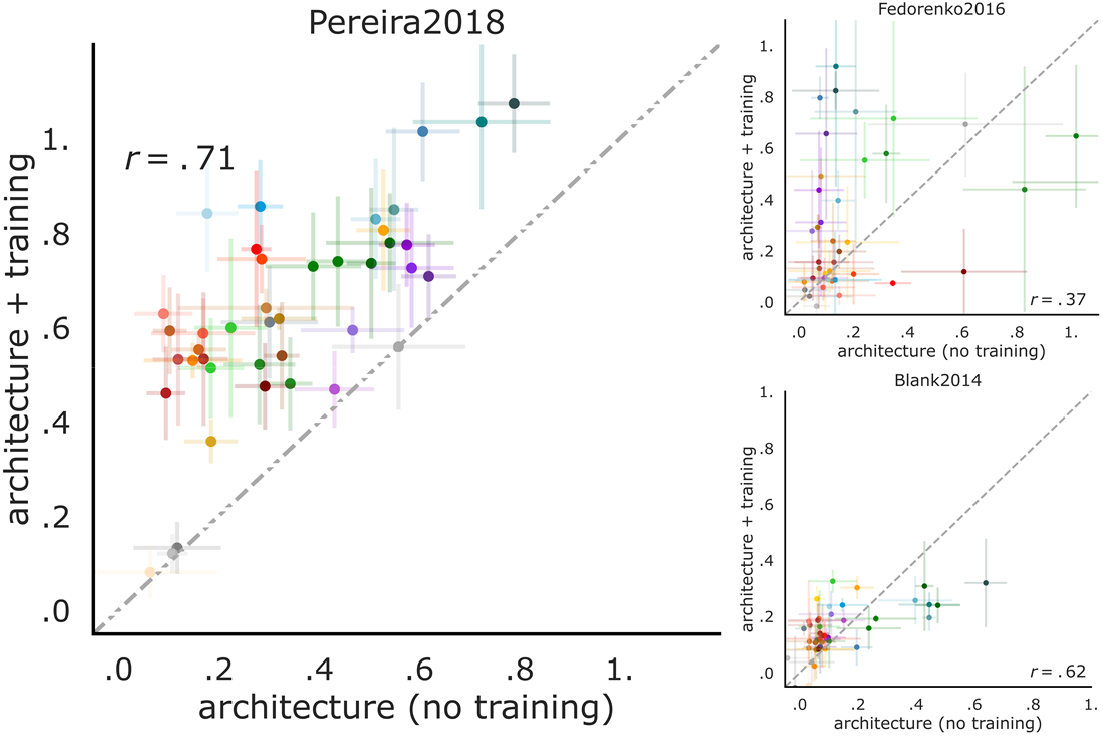
Model architecture contributes to brain predictivity and untrained performance predicts trained performance. In Fig. 5, we showed that untrained models already achieve robust brain predictivity (averaged across the three neural and one behavioral datasets). Here, we show that this relationship also holds for each dataset individually: Pereira2018: p<<0.00001, Fedorenko2016: p<0.05, Blank2014: p<0.00001.

**Figure S9:**
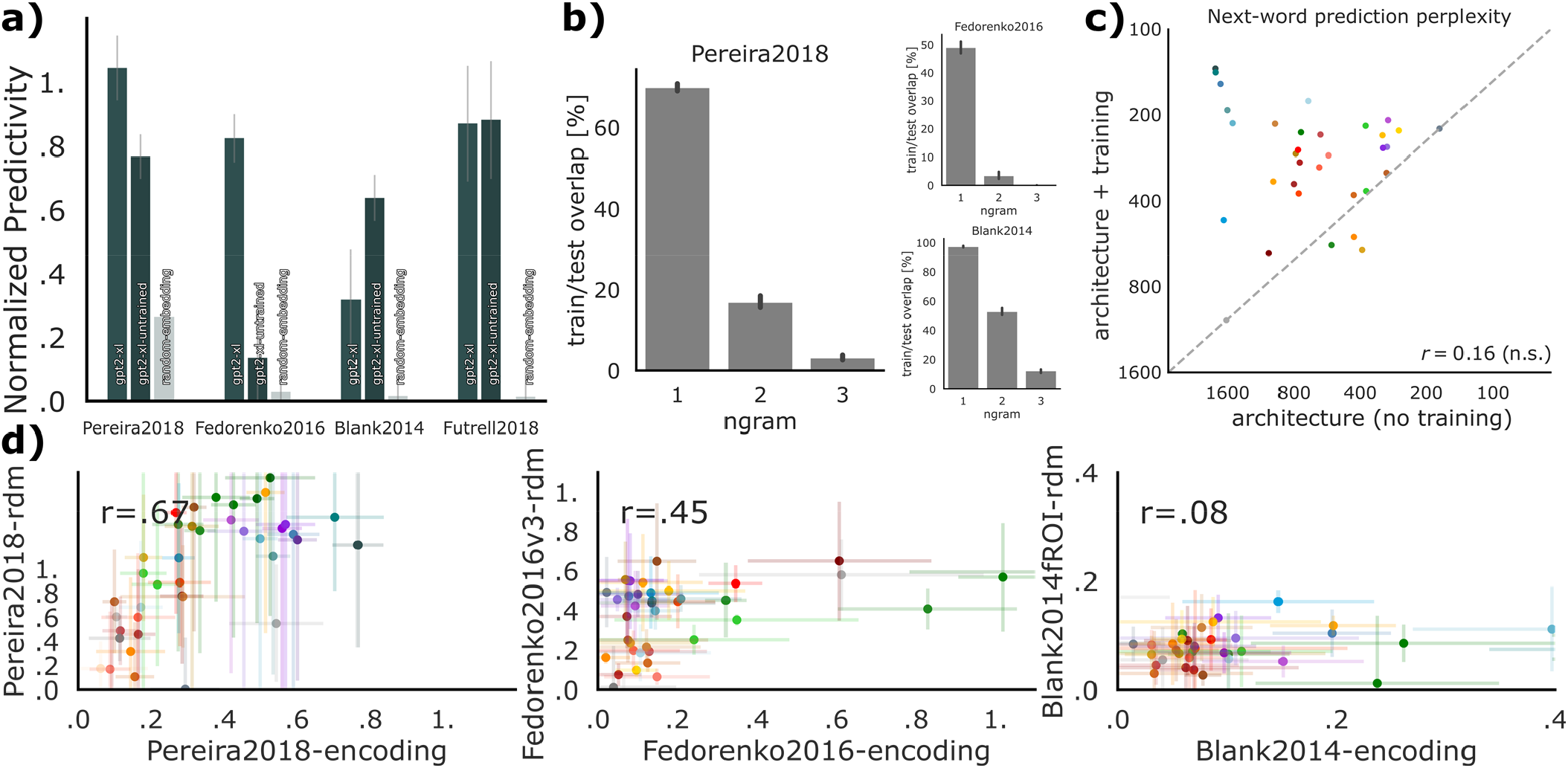
Controls for untrained models. **a)** Neural and behavioral scores of GPT2-xl, the best-performing model, with vs. without training, and of a random embedding of the same size. A large feature size alone is not sufficient: a random embedding matched in size to GPT2-xl scores worse than untrained GPT2-xl in all four datasets (3 neural, and 1 behavioral). These results suggest that model architecture critically contributes to model-to-brain and model-to-behavior fits. **b)** Overlap of bi- and tri-grams in train/test stimuli splits of benchmarks is minimal, and despite single-word overlap memorization of per-word responses is insufficient (a). **c)** The relationship between model performance with vs. without training on the wikitext-2 next-word-prediction task. Consistent with model performance with vs. without training on neural and behavioral datasets (Fig. 5), untrained models perform reasonably well. Training improves scores by 80% on average, and most prominently for GPT models, in teal (where the quality of the training data is optimized; see <u>Computational models</u> in <u>Methods</u>). GPT’s poor performance on next-word prediction might be explained by very high representational similarities across words pre-training in its last layer (Ethayarajh, 2019). **d)** Scores for untrained models obtained via linear predictivity generalize to scores obtained via RDM correlations. The RDM metric does not use any fitting. Correlations for untrained models’ scores between the predictivity and the RDM metric are: Pereira2018 r=.67, p<0.000005; Fedorenko2016 r=.45, p<.005; Blank2014 r=.08, n.s. See Fig. S2 for details on the RDM metric.

### SI-10 – Effects of model architecture and training on neural and behavioral scores

The 43 language models included in the current study span three major types of architecture: embedding models, recurrent models, and attention-based transformer architectures. However, in addition to this coarse distinction, the individual models vary widely in diverse architectural and training features. A rigorous examination of the effects of different model features on model-to-brain/behavior fit would require careful pairwise comparisons of minimally different models, which is not possible for ‘off-the-shelf’ models without extremely expensive re-training from scratch under many/all possible combinations of architecture, training diet, optimization objective, and other hyper-parameters. However, we here undertook a preliminary exploratory investigation. In particular, for a subset of model features (Table SI-9), we computed a Pearson correlation between the feature values and the averaged model score across all four datasets (3 neural, and 1 behavioral). We included five architectural features. Three features were continuous: i) number of hidden layers, which varied between 1 and 48 (mean 16.02, std. dev. 11.02); ii) number of features (units across considered layers), which varied between 300 and 78,400 (mean 20,971.26, std. dev. 18,362.91); and iii) the size of the embedding layer, which varied between 128 and 48,000 (mean 872.28, std. dev. 744.33). And the remaining two features were binary: iv) uni-vs. bi-directionality (32/43 models were bi-directional), and v) the presence of recurrence (5/43 models had recurrence). And we included two training-related features: i) training data size (in GB), which varied between 0.2 and 336 (mean 351.06 std. dev. 726.81); and ii) vocabulary size, which varied between 30,000 and 3,000,000 (mean 223,096.95 std. dev. 561,737.36). All training data numbers were taken from the original model papers, and if training data was specified in tokens, a conversion rate of 4 bytes per token was used. We further excluded the multilingual XLM and BERT models when examining the effect of training data size, because those numbers could not be confidently verified. For comparison, we also included performance on the next-word-prediction task that we examined in the main text.

The results are shown in Fig. S10. As expected—given the results reported in the main text for the individual datasets (Fig. 3, 4c)—next-word prediction performance robustly predicts model-to-brain/behavior fit (r = 0.49, p < 0.01). These results suggest that optimizing for predictive representations may be a critical shared feature of biological and artificial neural networks for language. How do architectural and training-related features compare to next-word-prediction task performance in their effect on neural/behavioral predictivity? Two architectural size features are most correlated with model performance: number of hidden layers (r = 0.56, p < 0.001), and number of features (r = 0.68, p << 0.0001). This is expected given that the most recent models with the highest performance on linguistic tasks are also the largest ones that researchers are able to run on modern hardware. The two training-related features—training data size and vocabulary size—are significantly *negatively* correlated with model performance. To rule out the possibility that the negative effect of training-related features is driven by models with relatively small training datasets and vocabulary size (e.g., ETM; Table S11) that have low brain/behavior predictivity, we ran an additional analysis considering only transformer models (n=38): even in these generally highly predictive models, more training data (r = -0.29, p = 0.11 [not plotted]) or larger vocabulary size (r = -0.21, p = 0.25 [not plotted]) do not appear to be beneficial, although the negative correlations are non-significant.

Does the collection of model designs investigated in this paper inform the hyperparameters that should be optimized for in any new model to achieve high predictivity? To provide a preliminary answer to this question, we performed an exploratory analysis in the form of stepwise forward model selection and examined (a) the most parsimonious model that explains the data, and (b) how much variance the selected features explain cumulatively (Fig. S10b). High overall explained variance indicates that the combination of features selected by the model is predictive of model performance, whereas low overall explained variance indicates that crucial predictive hyperparameters are still being neglected. In the forward regression analysis, we add predictors based on the highest R2-adjusted value of the new model, as long as variance increases by adding a new factor. This analysis revealed that adding training dataset size and recurrence does not lead to variance increase. Significance markers indicate the p-value for significance of adding each term, and for each regression step we plot the added explained variance (in R2-adjusted) of the variable chosen by the model. The overall cumulative R2-adjusted value of the selected model is 0.822.

**Figure S10:**
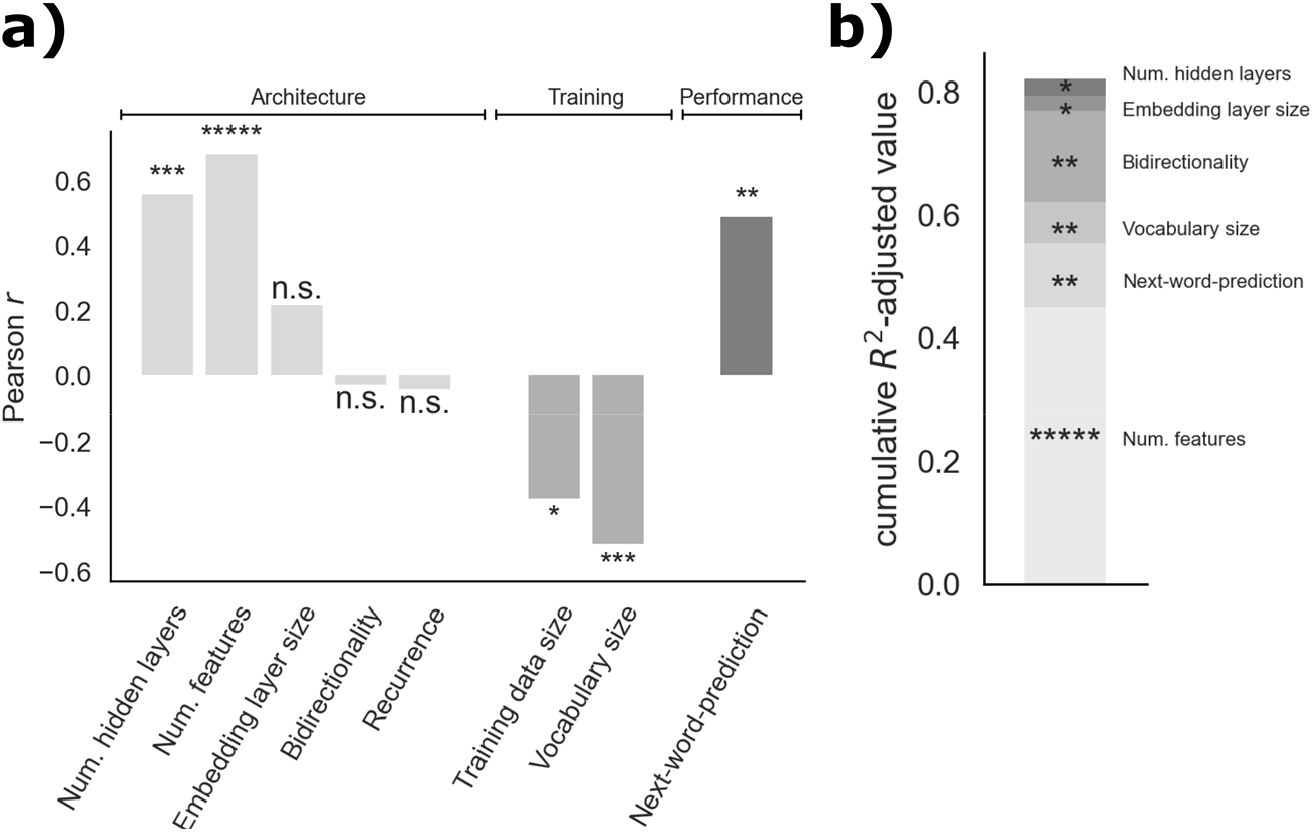
Effects of model architecture vs. training on neural and behavioral scores. **a)** We compared the effects on neural and behavioral scores (the averaged model score across all four datasets) of three kinds of features: (i) architectural properties, (ii) training-dependent variables, and, for comparison, iii) performance on the next-word-prediction task examined in the main text (Fig. 3, 4c). **b**) Alternative combination of predictors with stepwise forward regression model. New predictors are added based on the highest R2-adjusted value of the new model, as long as variance increases by adding a new factor (thus excluding training dataset size and recurrence). Significance markers indicate the p-value for significance of adding model terms. For each regression step, we plot the added explained variance (in R2-adjusted) of the variable chosen by the model. The overall cumulative R2-adjusted value of the selected model is 0.822. As in a), the preferred explanatory variable is the number of features. Stepwise forward regression based on significance leads to the same model-choice. Note that, as above, t5-11b is excluded for regression based on next-word-prediction, and multilingual models are excluded for regression on training size.

**Table S11:**
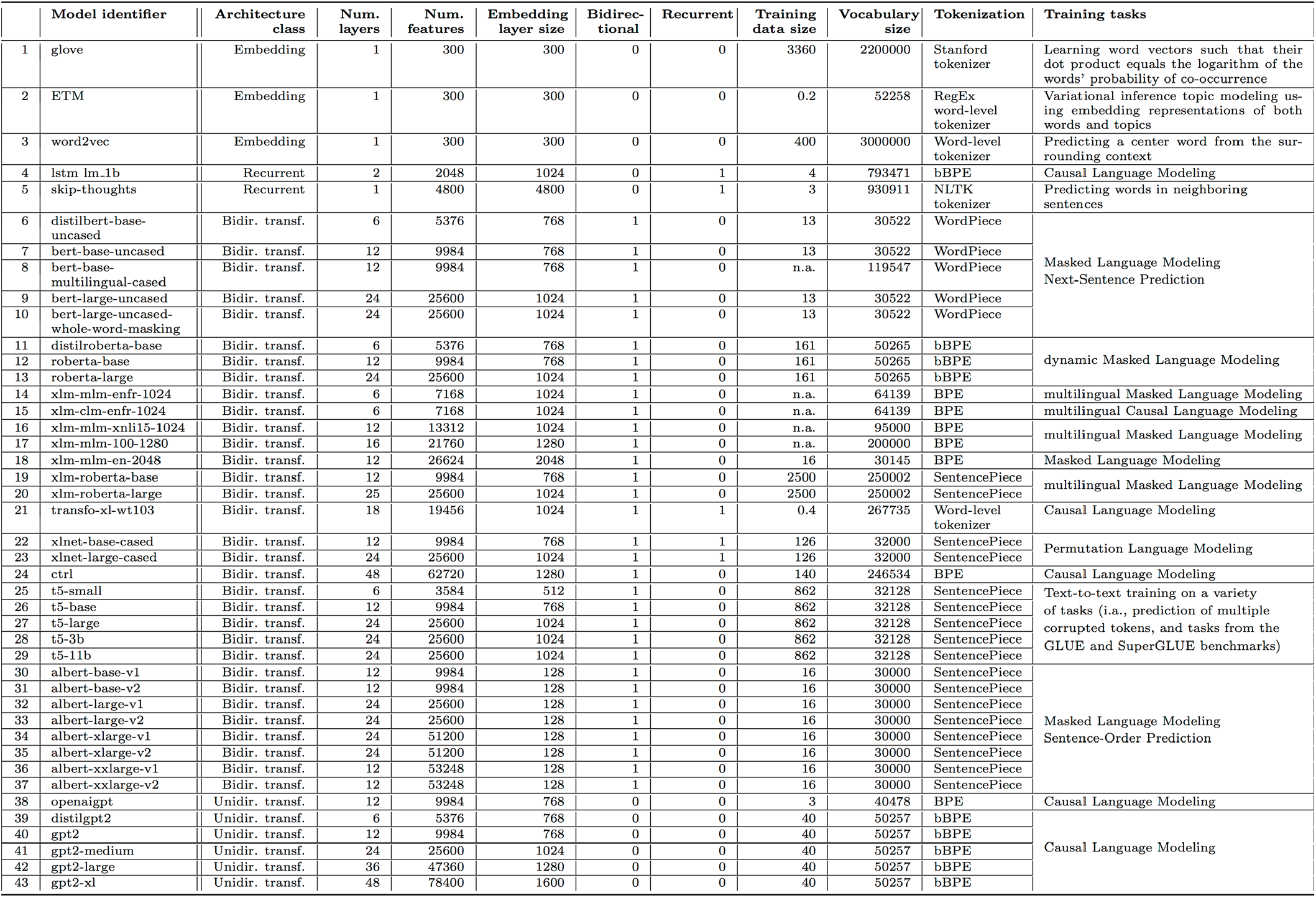
Overview of model designs.

**Figure S12:**
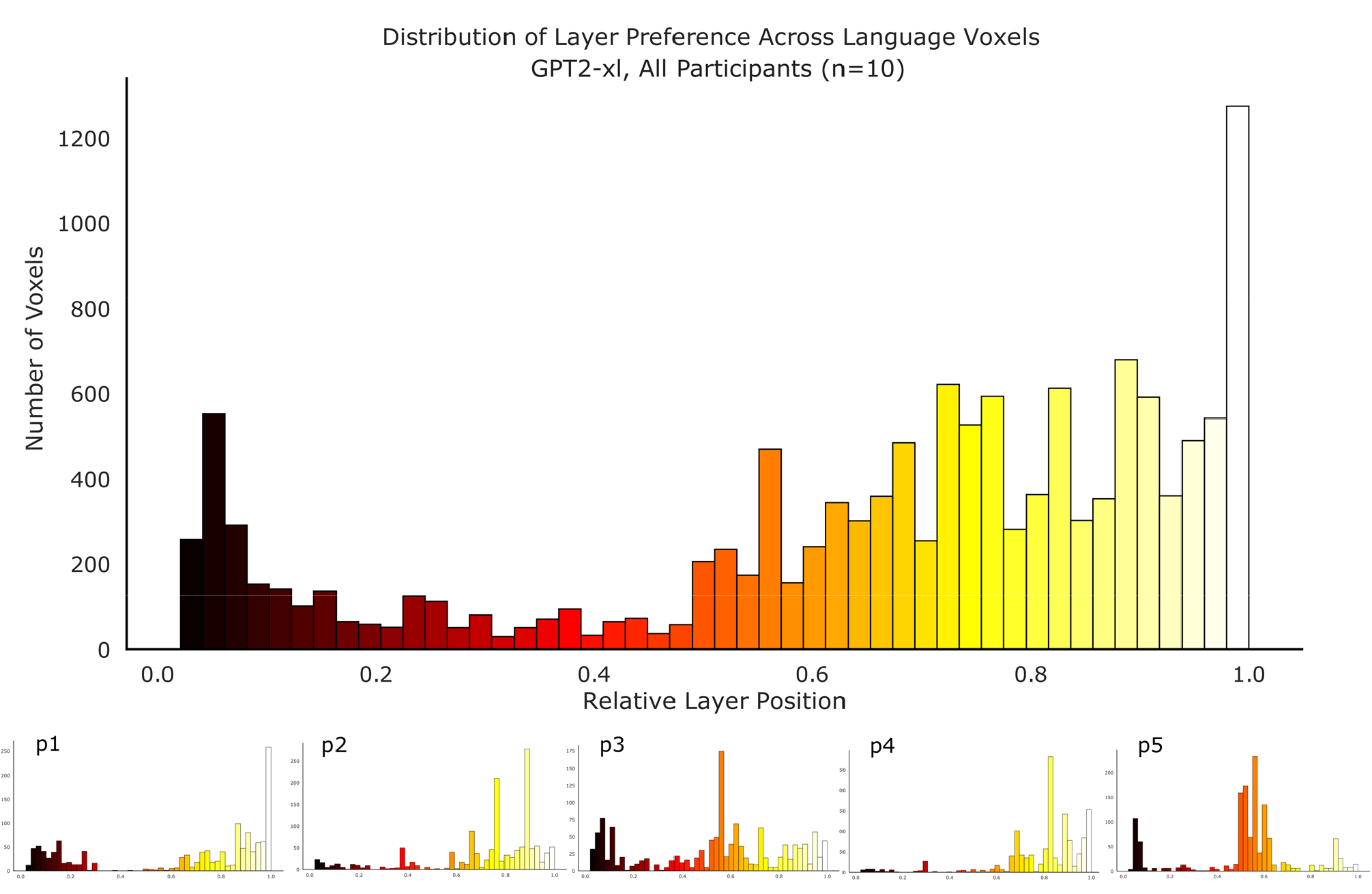
Distribution of layer preference (best performing layer) per voxel for GPT2-xl for *Pereira2018*. A per-voxel per-participant raw predictivity value (as opposed to *overall* ceiled predictivity scores in Fig. 2c) was obtained in the language network by computing the mean over cross-validation splits and experiments. For each voxel, the layer with the highest predictivity value was estimated as the “preferred” layer (argmax over layer scores). As in the main analyses, the voxels in the language network were included. Zero on the x-axis corresponds to the embedding layer of the model. The upper plot is averaged across all participants in *Pereira2018* (n=10). The lower panel shows the participant-wise layer preference for five representative participants. Across participants, most voxels show the highest predictivity value for later layers of GPT2-xl. Within participants, the layer preference across voxels varies but is often clustered around particular layers. Investigations of how predictivity fluctuates across model layers, and/or between the language network and other parts of the brain, is left for future work.

**Figure S13:**
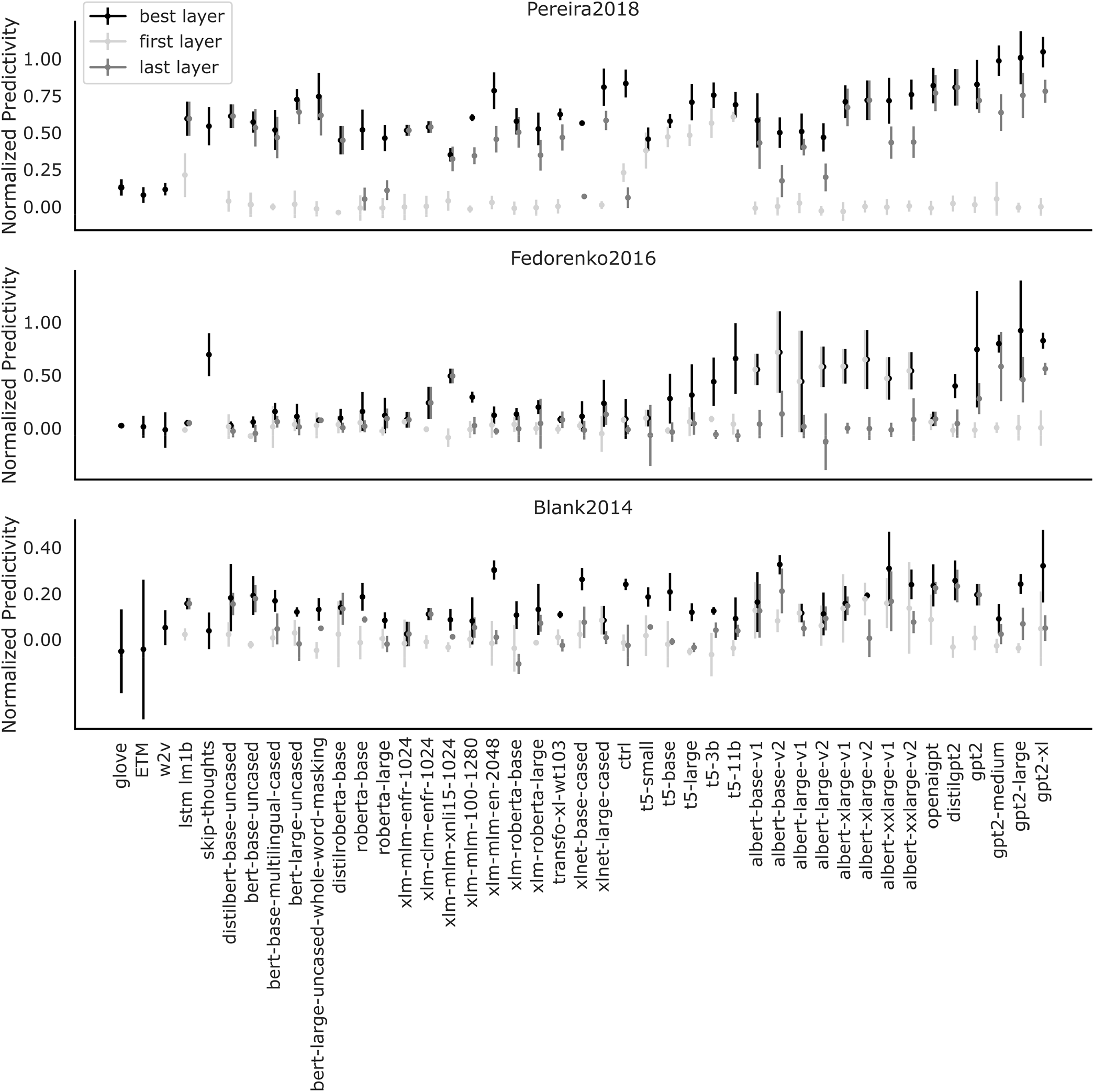
Brain scores of each model’s best, first, and last layer. To test the importance of intermediate representations, we directly compared layer performances at the beginning and end of each model with the model’s best-performing layer. In nearly all networks with multiple layers, both the token embedding (first layer) as well as the task-specific output (last layer) underperform significantly compared to the respective best layer. This suggests that the combination of architecture and weights in the networks is a major driver for brain-like representations, beyond potential semantic information that is already present in the model input codes. Lexical similarity determined by optimizing for next-word prediction present in the output layer is also not sufficient, instead pointing to intermediate representations as the most predictive (see also Fig. 2c).

